# Butterfly egg plasticity in physiologically and reproductive relevant traits: adaptation to climate?

**DOI:** 10.1101/2025.08.29.673100

**Authors:** Teun de Jong, Liana O. Greenberg, Ella Catley, Nina Fatouros, Graziella Iossa

## Abstract

Embryos of terrestrial animals developing inside immobile eggs face variable environmental conditions. Insect egg pores, critical for gas exchange (aeropyles) and fertilisation (micropyles), are ecologically important yet understudied. While aeropyles likely adapt to climatic respiratory demands, and micropyles may respond to environmental pressures, their inter- and intra-specific variation remains poorly studied. Here, we compared egg pore morphology of three closely related *Pieris* butterflies sampled in the Netherlands. In addition, we compared egg traits of different *P. napi* populations sampled along a 4,000-km latitudinal gradient from northern to southern Europe linked these traits with available climatic data. After several generations under identical greenhouse conditions, egg pore traits remained highly variable within and between species and groups. Remarkably, *P. napi* populations from divergent climates differed more in pore number and size than did co-occurring *Pieris* species. Aeropyle number and width were strongly associated with climatic variables such as maximum temperature, independent of egg size. Micropyle width varied with seasonality. Additionally, we observed the micropylar pit of older eggs covered by a substance more often than fresh eggs. Our findings indicate that insect egg pores are highly dynamic traits, shaped by ecological, behavioural and climatic factors and an under-recognised layer of egg adaptation.

## Introduction

In most insects, parental care is limited to the choice of oviposition site, after which egg survival depends on egg traits^1,2^. Being immobile and unable to relocate to preferred conditions, eggs represent a vulnerable life stage for insects^2–4^. Additionally, insect eggs are tiny, with a large surface area relative to volume, making the thin eggshell a critical mediator between the embryo and its external environment^5,6^. The insect eggshell is typically waterproof, resistant to UV radiation and desiccation, while balancing the competing demands of respiration and water conservation^5^. For many insects, respiration and fertilisation are mediated by minute pores on the outer eggshell, known as aeropyles and micropyles, respectively^5,7^. Although these pores are essential for embryonic development, they remain relatively understudied and somewhat overlooked.

While it is well established that environmental variation drives genetic and phenotypic adaptation in many insect egg traits^8^, the extent to which this applies to egg pores remains poorly understood^9^. Despite the large interspecific diversity in pore morphology^6,9^, most literature has focused on pore morphogenesis^7,10–12^, their role in pesticide permeability^13–15^, or their utility for taxonomic classification^16,17^, rather than the evolutionary pressures shaping their diversification^18^. It has been hypothesised that the remarkable diversity in egg pore structure reflects functional divergence related to adaptation to environmental conditions, thereby mediating embryo development^5,6,19^. More broadly, the drivers of egg morphological diversification, such as egg size, have recently received renewed attention^20,21^. However, only a handful of studies have investigated how insect egg pore traits respond to abiotic stresses or divergent local condition^6,9,19,22^.

Aeropyles are specialised structures that allow airflow between the external environment and the spongy chambers of the inner exochorion^5^. In many species, covering the aeropyles with a non-permeable substance causes embryo mortality, highlighting their critical role in respiration^5,22^. All terrestrial insects and their eggs face a trade-off between permitting gas exchange to facilitate metabolic processes, and preventing water loss^23^. This trade-off is highly temperature-dependent because, at higher temperatures, development is faster and metabolic demands increase. Therefore, embryonic oxygen requirements increase, but the rate of water loss and risk of desiccation are also higher^24^. Insufficient oxygen exchange may lead to anoxic regions in the developing embryo^5,23,25^. Thus, the number and size of aeropyles likely play a role in regulating this trade-off. For example, a smaller total aeropyle area (either by fewer or smaller aeropyles or both) may conserve water but limit oxygen availability. In contrast, greater aeration (either by more numerous or larger aeropyles or both) may facilitate rapid growth but increase the risk of desiccation.

Egg size is another factor that may influence the trade-off between respiration and water conservation during embryonic development^26,27^, and it, too, varies with latitude and climate. Larger eggs typically contain larger embryos, which require more oxygen^26,27^, and egg size often increases with latitude in several insect taxa^27–30^. This trend aligns with Bergmann’s rule, formulated initially for endotherms^31^, which has been observed in many, but not all, insect species along latitudinal gradients^32–34^. An alternative, the “canteen hypothesis,” proposes that larger eggs buffer against desiccation in hot, dry climates by storing more water^26,35^. However, empirical support is mixed, and other factors such as clutch size may better explain variation in desiccation risk. Grouped eggs tend to lose less water than solitary ones^27,36,37^, and clutch size often correlates with egg size^27,28,34^. Recent large-scale analyses of insect eggs suggest that ecological factors, particularly oviposition environment, better explain variation in egg morphology than do universal allometric or physiological rules^38^. Consistent with the finding that ecological factors explain egg morphology, we hypothesise that aeropyle area, rather than egg size per se, shows climate-linked variation. To clarify the role of egg size and whether selection on egg size interacts with the evolution of aeropyles and micropyles^20,26,28^ we require studies that simultaneously examine both egg size, egg number, and egg pore traits across diverse climates^20,26,28^.

Micropyles are narrow channels that traverse both the exo- and endochorion of the insect eggshell and function as conduits through which sperm reaches the vitelline membrane during fertilisation^5,10,11,14.^. Although micropyle number varies widely among insect species, it is typically conserved within species^9^. These structures may also mediate post-copulatory sexual selection, including sperm competition and cryptic female choice, where multiple micropyles allow the chance for fertilisation by multiple sperm^39^. Phylogenetic studies further suggest a relationship between micropyle number and the degree of polyandry or mating frequency across species, with promiscuous species having more micropyles^9^. Despite this knowledge, drivers of divergence in micropyle size between or within species have not yet been investigated. Beyond their role in fertilisation, micropyles may also contribute to embryo survival in ecologically meaningful ways. A recent phylogenetic analysis found that micropyle number is positively correlated with both aeropyles and annual precipitation: species from drier climates tend to have fewer micropyles and aeropyles, potentially reducing egg vulnerability to desiccation^40^. Gas exchange can indeed occur through micropyles^41,42^. We expect to find differences in micropyle traits both within and between species, which are associated with divergent egg-laying strategies and climates. In addition to the adaptive features of micropyles, a cover may protect the egg from environmental stress by coating the micropyle. For instance, in large cabbage whites (*Pieris brassicae*), the yolk retracts from the anterior pole post-oviposition, and an oily layer covers the micropyles^14^. Since this oily material can prevent water loss^43,44^, and gas exchange can occur through micropyles^41,42^, sealing the micropylar pit may represent an important adaptive mechanism against desiccation. Very few studies have documented or described these egg-derived coating substances in butterfly eggs^14^. We expect to find differences in micropyle traits both within and between species, which are associated with divergent egg-laying strategies and climates. In addition to the adaptive features of micropyles, a cover may protect the egg from environmental stress by coating the micropyle. For instance, in large cabbage whites (*Pieris brassicae*), the yolk retracts from the anterior pole post-oviposition, and an oily layer covers the micropyles^14^. Since this oily material can prevent water loss^43,44^, and gas exchange can occur through micropyles^41,42^, sealing the micropylar pit may represent an important adaptive mechanism against desiccation. Very few studies have documented or described these egg-derived coating substances in butterfly eggs^14^.

Here, we use a common garden approach to investigate variation in egg pore traits in cabbage white butterflies (*Pieris* spp.), focusing on both interspecific and intraspecific patterns. We assess (i) interspecific differences in egg size, aeropyles, and micropyles among three co-occurring species (*P. napi, P. rapae, P. brassicae*); (ii) intraspecific variation in these traits across a 4,000-km climatic gradient in *P. napi*; and (iii) post-oviposition changes in micropylar coverage. We hypothesise that aeropyle and micropyle traits vary with local climate factors. Specifically, we predict that egg pore traits, more than egg size, reflect bioclimatic pressures. Consistent with this, we find that pore morphology diverges most strongly in *P. napi* populations at climatic extremes, supporting a role for local adaptation.

## Materials and methods

### Insects and plants

*Pieris spp*. butterflies (Lepidoptera: Pieridae) are common all over Europe and can be found where their crucifer (Brassicales: Brassicaceae) host plants occur. Species differ in their preferred microclimate selected for oviposition, with *P. brassicae* and *P. rapae* generally ovipositing in similar conditions, often in drier habitats, including open fields^45,46^. *Pieris napi* instead selects a moist habitat, often near water when in drier environments^45,47^. Females of *P. napi* and *P. rapae* lay eggs singly, while *P. brassicae* females lay eggs in clutches^47^. These three species all prefer brassicaceous leaf tissue for oviposition. Their eggs, less than 1.5 mm in length, are laid on the lower side of leaves, where the leaf boundary layer provides a moist microclimate and a buffer from high temperatures^2,48,49^.

To explore interspecific diversity, we analysed 14 egg clutches from *P. brassicae* (in total 69 eggs), 17 eggs from *P. rapae*, and 25 eggs from *P. napi*. Butterflies were lab-reared and originated from the surroundings of Wageningen, the Netherlands (51°97’91.19”N, 5°68’52.59”E). Butterflies were first collected in 2020 and the rearings were annually refreshed. All species were reared on black mustard (*Brassica nigra*) and/or Brussels sprouts plants (*Brassica oleracea* var. *gemmifera*) in greenhouses (Unifarm) at Wageningen University (18–20°C, 50–70% RH, photoperiod L16: D6). Individual butterfly populations were separately caged and comprised 20-30 adults, fed with a 20% honey/water solution, and allowed to mate freely.

To explore intraspecific diversity in *P. napi* egg traits, we used six additional populations, wild-caught in 2022 and 2023 across Europe. Butterflies were reared for a minimum of three to a maximum of ten generations under greenhouse conditions. From northernmost to south, we analysed eggs from populations originating from (i) Abisko, Sweden (68°34’76.10”N, 18°82’97.70”E), 25 eggs; (ii) Stockholm, Sweden (59°36’44.70”N, 18°03’11.50”E), 25 eggs; (iii) Wageningen, the Netherlands (51°97’91.9”N, 5°68’52.59”E), 25 eggs; (iv) Jura, France (46°55’57.7”N, 6°08’49.60”E), 22 eggs; (v) Callas, France (43°59’03.0”N, 6°53’12.66”E), 25 eggs; (vi) Pyrenees, Spain (42°32’28.3”N, 1°06’42.98”E), 13 eggs; and (vii) Costa brava, Spain (42°26’49.5”N, 3°06’90.01”E), 20 eggs (Figure 1*b*). In the wild, the populations from Abisko and Stockholm are univoltine and bivoltine, respectively. Under greenhouse conditions, the butterflies naturally enter diapause after a few generations. We considered a pupa to be diapausing if it did not emerge within a month. These pupae were transferred to a refrigerator (4℃) for three to six months and then returned to greenhouse conditions for emergence. Therefore, the number of generations kept in captivity differed between populations. For cryo-scanning electron microscopy, eggs still in position on their host leaf were obtained from adult butterflies kept in the greenhouse and ovipositing freely on black mustard (*Brassica nigra*) plants in February 2023 and April 2024.

**Figure 1.**
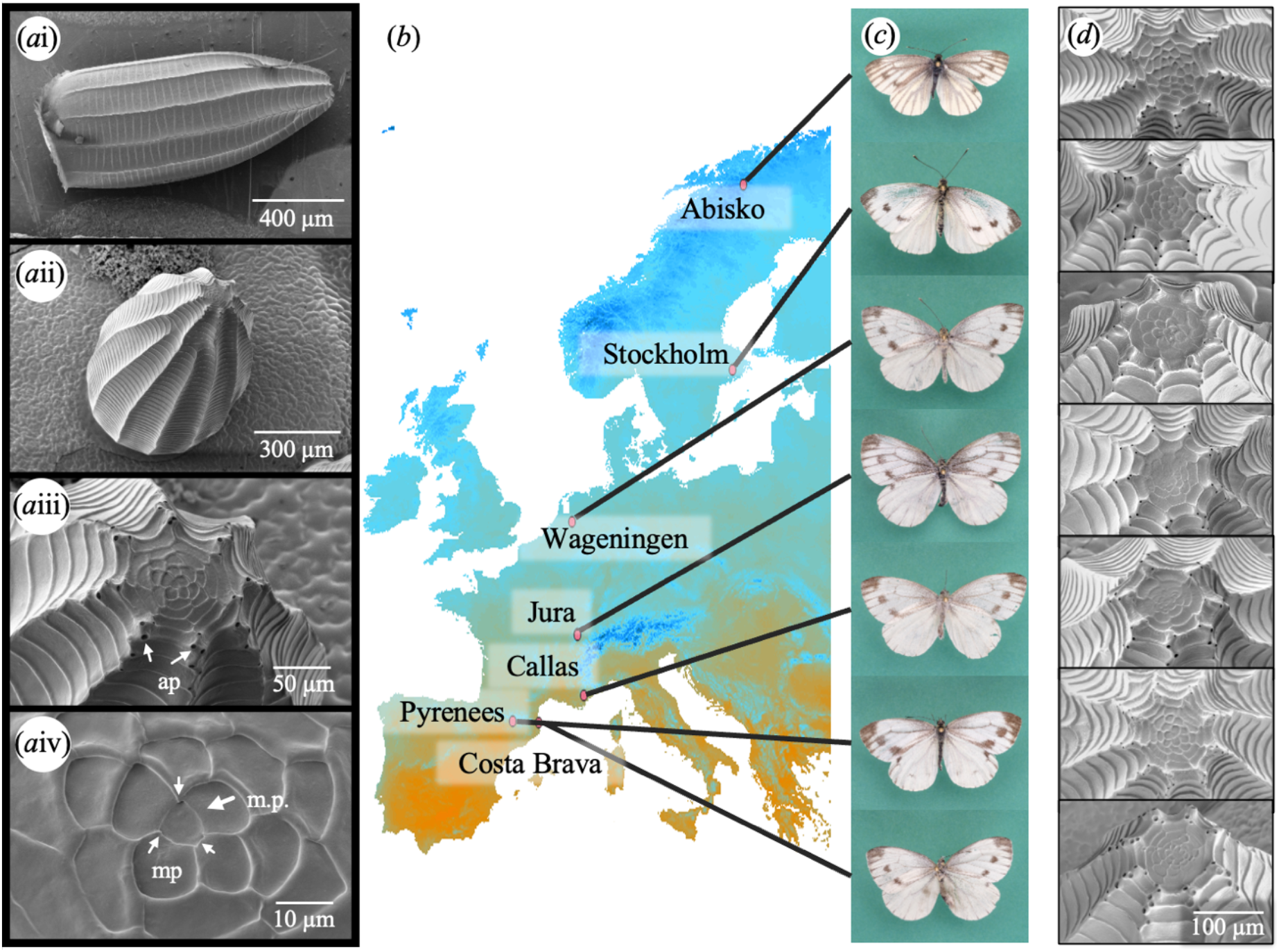
*Pieris napi* populations collected from North to South Europe and their egg and butterfly wing morphology. (*a*i-iv) Cryo-scanning electron microscopy images (cryo-SEM) of egg shapes and pores taken from a Stockholm *P. napi* egg. (*a*i) Whole egg image, showing full length. (*a*ii) Whole egg image, showing the abaxial leaf surface and the apical side of the egg. (*a*iii) Closeup of egg tip showing the aeropyles (ap; arrows). (*a*iv) Micropylar pit (m.p.), showing the openings of three micropyles (mp; small arrows). (*b*) Map showing the origin of butterfly populations. Colours indicate the maximum temperature of the warmest month, with orange representing high and blue representing low temperatures. (*c*) Variety in butterfly wing melanisation across the seven *P. napi* populations. (*d*) The SEM images from the different populations with the egg ridges leading up to the apical egg tip, where the egg pores are located. Aeropyles are located on ridges, and micropyles are in the centre of the eggtip (not visible).

### Cryo-scanning electron microscopy

We imaged a total of 241 eggs of *Pieris* spp. containing 427 micropyles and 10195 aeropyles. *Pieris* eggs of one to three days of age, still in position on the *Brassica nigra* leaf they were laid on, were attached to a sample holder using a very thin layer of Carbon adhesive glue (Leit-C, EMS, Washington, PA, USA). Samples were frozen by plunging them into liquid nitrogen and subsequently placed in a cryo-preparation chamber (MED 020/VCT 100, Leica, Vienna, Austria). To sublimate any water vapour contamination (ice) from the surface, the samples were placed for 10 minutes at -92°C. Samples were then sputter coated with an 8 nm layer of tungsten from three angles and transferred under vacuum to the field emission scanning electron microscope (Magellan 400, FEI, Eindhoven, the Netherlands) onto the sample stage at -120°C. The images were taken with the secondary electron detector set at 2kv, 13pA. For each egg, a minimum of three cryo-scanning electron microscopy (cryo-SEM) images were captured; the whole egg for size measurements (Figure 1*a*ii), the apical egg tip to visualise aeropyles (Figure 1*a*iii), and the micropylar pit to measure micropyle number and width (Figure 1*a*iv). We used general tools from the FIJI platform to measure egg pores on the cryo-SEM images^50^.

### Local climate data at field sites

The bioclimatic variables from WorldClim^51^ are widely used to study climatic effects on insects and other animals. We selected three variables: temperature seasonality (BIO4, the annual standard deviation of monthly mean temperatures), maximum temperature (BIO5, the average maximum temperature of the warmest annual month), and annual precipitation (BIO12). These variables are of biological significance: temperature and humidity directly influence the trade-off between insect gas exchange and water loss^23^, and precipitation was previously found to correlate with micropyle number in insect eggs^40^. Temperature seasonality reflects the variation of temperature across an insect’s lifetime and could thus directly (e.g. by a relatively cold start of the season) or indirectly (via other life stages) affect insect egg development^52,53^. Furthermore, we selected the length of the insect growing period, as this affects generation length and, thus, potentially constrains the developmental time of eggs and larvae. Data on the length of the growing period is referred to as ‘growing days’, which is the number of days when the mean daily temperature exceeded 10°C. This temperature is often used as the minimum threshold for growing degree days in Lepidoptera ^54.^ These data were obtained from the public dataset of the Food and Agriculture Organization of the United Nations^55^ (available at https://gaez.fao.org/).

The freely accessible bioclimatic variables from the WorldClim dataset span 1970-2000. To account for recent changes in the local climate, we calculated the same variables for a more recent timespan, 2018-2021, using the ‘biovars’ function of the R package *dismo*^*56*^. This package requires monthly values for precipitation and minimum and maximum temperatures, which we extracted from the 2.5 minutes spatial resolution data from the WorldClim historical monthly weather dataset CRU-TS 4.06^57^, downscaled with WorldClim 2.1^51^, at our field site coordinates (Table S3).

### Statistical analyses

To test for phenotypic adaptation, we analysed five different egg characteristics: egg width, micropyle number, micropyle width, aeropyle number, aeropyle width, and presence/absence of a micropylar pit cover. We found egg width to be a suitable proxy for egg size (Figure S1, Table S1). For each trait, we constructed either a linear model (*lm*) using base functions in *R*^58^or a mixed effect model (*lmer*) using the *R* package *lme4* when accounting for random effects^59^. As there are multiple aeropyles and micropyles per egg, we treated each measured pore width as an individual data point and included egg ID as a random effect. *Pieris brassicae* butterflies lay eggs in clutches. Therefore, we imaged five eggs per clutch and included clutch ID as a random effect in the models for this species. We separately analysed interspecific differences (three species obtained from Wageningen, n = 46 eggs/clutches) and intraspecific differences (seven populations of *P. napi*, n = 156 eggs). We could not measure all traits from all eggs (e.g. due to a micropylar cover or due to the cryo-SEM angle; Table S2).

We tested the model fits using the *DHARMa* package^60^. To analyse categorical data, we compared nested models with the *R* base function ‘Anova’. We performed pairwise comparisons using estimated marginal means (EMMs) obtained using the ‘emmeans’ function from the *emmeans* package with the Tukey method for p-value adjustment^61^. We tested whether *P. napi*’s egg width was associated with aeropyle number, aeropyle width, micropyle number, and micropyle width. Therefore, we did not correct for egg width in our comparative models. We used eggs from all species and populations to analyse the increase in micropylar cover with egg age, and constructed a general linear model (*glm*) with the *binomial* family using base functions in R^58^ Here, we excluded 14 eggs that could not be assigned to an age group (remaining n = 217) and included both species and population as random effects.

### Structural equation model construction

We used a piecewise structural equation model (pSEM) to resolve the relationship between egg pore traits and egg size using the *R* package *piecewiseSEM*^*62*^. In the model, we included four climatic variables: temperature seasonality, the maximum temperature of the warmest month, annual precipitation, and growing degree days (n = 7 locations). Furthermore, we included five egg variables: egg width (n = 152 eggs), micropyle number (n = 64 eggs), individual micropyle width (n = 165 micropyles), aeropyle number (n = 94 eggs), and individual aeropyle width (n = 4361 aeropyles). Plotting the data indicated the possibility of a polynomial relationship between aeropyle traits and maximum temperature; therefore, we calculated the orthogonal polynomial of maximum temperature and included it as a variable (Supplementary 5). For model selection, we followed the procedure outlined in Boisseau and Woods 2024^20^ and described model selection in Supplementary 5. Briefly, the initial model encompassed all possible paths from climatic to egg variables and among egg variables. Then, the most parsimonious model was identified by comparing CIC values, dropping pathways one by one until the CIC value of the main model was lower than that of any of its reduced model versions. In the next step, we tested against the alternative direction of association between variables. To test the assumptions, all individual linear models were investigated using the *simulationResiduals* function of *R* package *DHARMa*^60^.

## Results

### *Pieris* egg characteristics

Eggs are shaped approximately as half an ellipsoid, with the anterior end being slightly extended and narrow (Figure 1*a*i, *a*ii). The posterior end is blunt and glued to the plant surface, usually on the abaxial surface of a leaf. Eggs are bright yellow to orange and typically contain 12 to 16 ridges, often extended along the complete length of the egg. Aeropyles are generally located on nodes on the anterior end of these ridges (Figure 1*a*iii). The ridges extend into a relatively flat crater. The micropylar pit, containing between one and six micropyles, is located roughly in the centre of this crater (Figure 1*a*iv).

### Interspecific differences

Species varied in their egg width (Anova: *X*^2^ = 99.7, df = 2, *p* < 0.001). Clustered *P. brassicae* eggs were significantly wider than the singly laid *P. rapae* (estimate = 113 ± 8 µm, *t* = 14.5, *p* < 0.001) and *P. napi* eggs (estimate = 112 ± 7µm, *t* = 15.9, *p* < 0.001), while *P. rapae* and *P. napi* were similar in width (estimate = 1 ± 8 µm, *t* = 0.1, *p* = 0.990; Table S2; Figure S2).

Aeropyle number did not significantly vary between species (Anova: *X*^2^ = 5.1, df = 2, *p* = 0.079; Figure 2*a*i), but aeropyle width did vary (Anova: *X*^2^ = 25.3, df = 2, *p* < 0.001; Figure 2*b*i). *Pieris napi* and *P. rapae* were similar in aeropyle number and width (*P. napi - P. rapae* aeropyle number: estimate = 0.39 ± 3.76; aeropyle width: estimate = -0.11 ± 0.18 µm, *t* = -0.60, *p* = 0.830). *Pieris brassicae* tended to have more aeropyles (n.s., see Anova results above), and they were significantly smaller (*P. brassicae-P. napi*: estimate = -0.74 ± 0.17 µm; *t* = -4.30, *p* < 0.001, *P. brassicae - P.rapae*: estimate = -0.85 ± 0.16 µm, *t* = -5.30, *p* < 0.001; Figure 2*b*i; Table S2).

**Figure 2.**
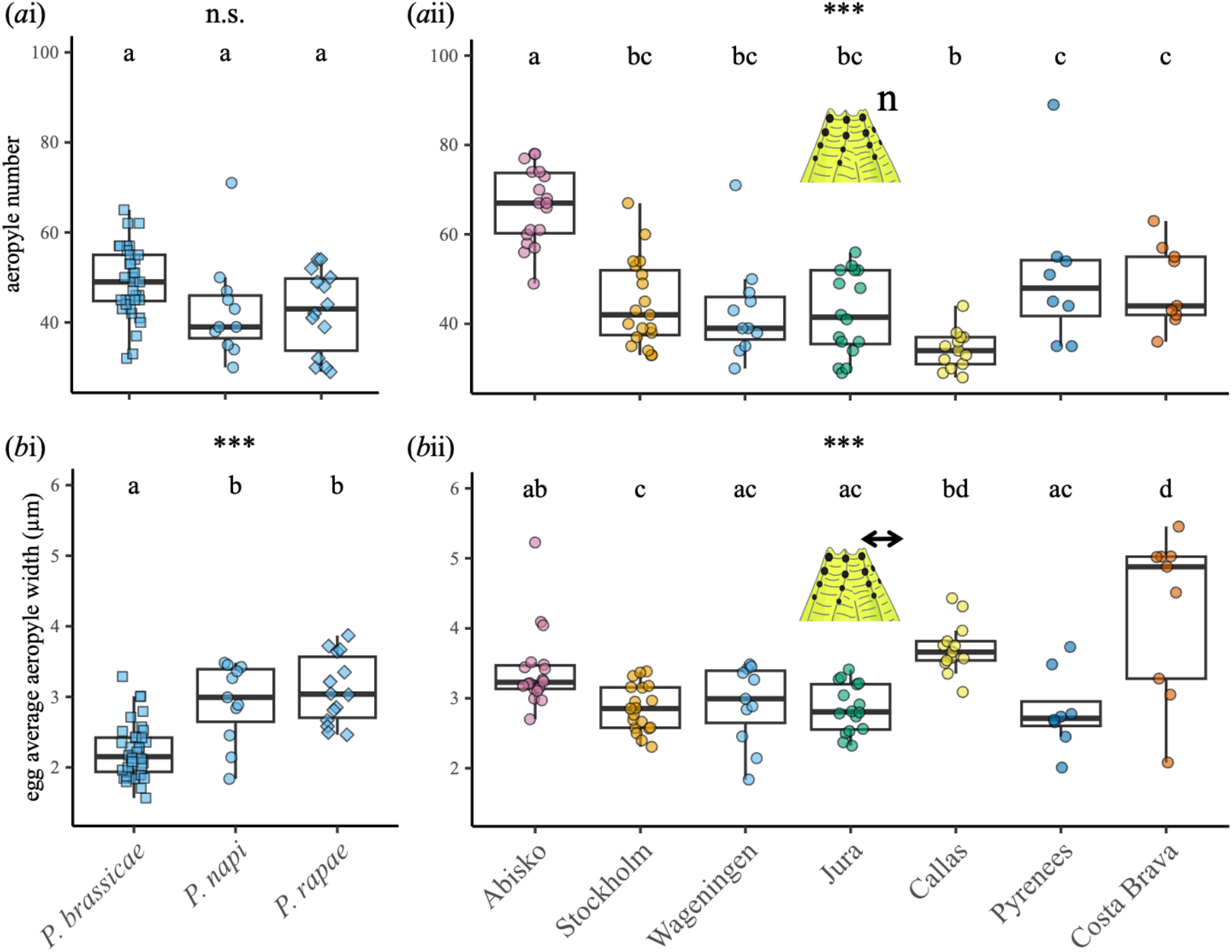
Comparison of egg aeropyle characteristics (*a*i, *b*i) interspecifically between *Pieris spp*. and (*a*ii, *b*ii) intraspecifically between *P. napi* populations. (*a*i, *b*i) Aeropyle number and average aeropyle width between species (n = 61 eggs and n = 2830 aeropyles, respectively). (*a*ii, *b*ii) Aeropyle number and average aeropyle width between populations (n = 94 eggs and n = 4411 aeropyles, respectively; Table S2; Table S5; Table S6). Plotted points represent individual eggs. Asterisks indicate the overall significance between compared groups, obtained with an Anova test (n.s. = not significant; *** *p* < 0.001). Different letters indicate significant differences (*post hoc* using EMMs, *p* > 0.05).

Species differed in micropyle number (Anova: *X*^2^ = 17.1, df = 2, *p* < 0.001; Figure 3*a*i), with *P. napi* having fewer micropyles (*P. napi - P. brassicae*: estimate = -1.19 ± 0.29, *t* = -4.10, *p* < 0.001; *P. napi - P. rapae*: estimate = -1.30 ± 0.40, *t* = -3.25, *p* = 0.005), and *P. rapae* having a similar number of micropyles as *P. brassicae* (estimate = 0.11 ± 0.36, *t* = -0.31, *p* = 0.950; Figure 3*a*i). The width of micropyles also significantly differed between species (Anova: *X*^2^ = 27.1, df = 2, *p* < 0.001; Figure 3*b*i), with *P. napi* having smaller micropyles (*P. napi - P. brassicae*: estimate = -0.46 ± 0.08 µm, *t* = 5.57, *p* < 0.001; for *P. napi - P. rapae*: estimate = -0.55 ± 0.14 µm, *t* = - 4.00, *p* < 0.001), while there was no significant difference between *P. brassicae* and *P. rapae* (estimate = -0.09 ± 0.13 µm, *t* = -0.64, *p* = 0.801; Figure 3*b*i; Table S2).

**Figure 3.**
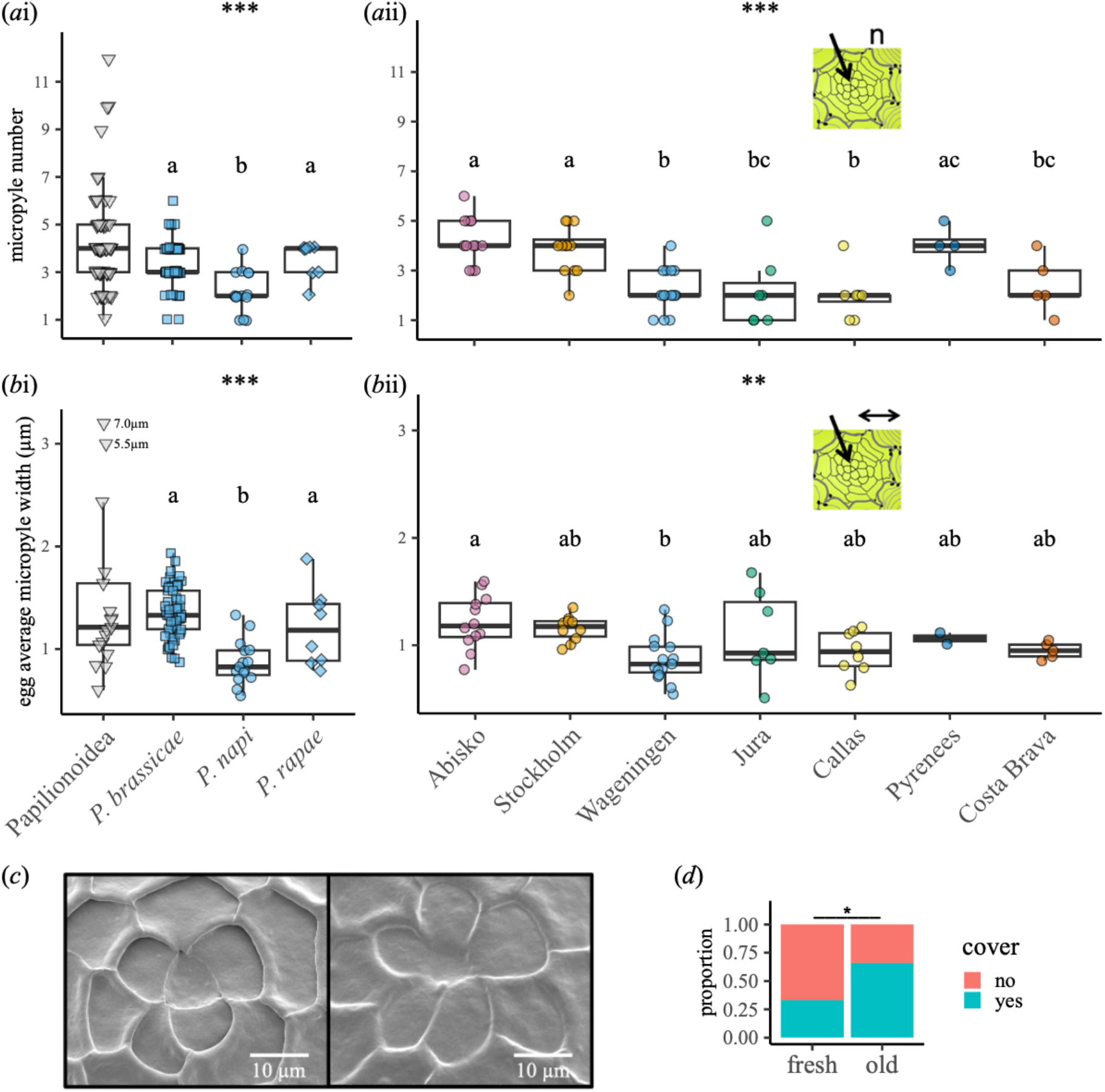
Comparison of egg micropyle characteristics (*a*i, *b*i) interspecifically between *Pieris spp*. and species of butterflies (Papilionoidea) and (*a*ii, *b*ii) intraspecifically between *P. napi* populations. (*a*i, *b*i) Micropyle number and average micropyle width between species (n = 81 eggs and n = 255 micropyles, respectively). (*a*ii, *b*ii) Micropyle number and average micropyle width between populations (n = 64 eggs and n = 165 micropyles, respectively; Table S2; Table S7; Table S8). (*c, d*) Coverage of the micropylar pit. (*c*) Cryo-SEM images show examples of an uncovered micropylar pit (left) and a fully covered micropylar pit (right). Plotted points represent individual eggs or averages of individual species for Papilionoidea. For the analysis of micropyle width, we used the values of individual egg pores. (*d*) Stacked bar plot showing the proportion of eggs of which the micropylar pit was covered to at least some extend for both fresh eggs (laid less than 2 hours before imaging) and older eggs (laid 4 hours or more before imaging; n total = 217 eggs). Data on Papilionoidea micropyles were obtained from^40^. Asterisks indicate the overall significance between compared groups (Papilionoidea was not included in comparisons), obtained with an Anova test (** *p* < 0.01; *** *p* < 0.001). Different letters indicate significant differences (*post hoc* using EMMs, *p* > 0.05).

### Intraspecific differences between *Pieris napi* between populations

Egg width, used as a proxy for egg size, significantly differed between the populations of *P. napi* (Anova: *F* = 20.8, *p* < 0.001; Table S2; Figure S2). Eggs from Abisko tended to be larger than all other populations. Mainly, eggs from Wageningen, Jura, Callas, and Costa Brava were significantly smaller (Table S2; Figure S2; Table S9).

Between the seven populations, there was a significant difference between both the aeropyle number (Anova: *F* = 16.57, *p* < 0.001; Figure 2*a*ii; Table S2; Table S5) and the individual aeropyle width (Anova: *X*^2^ = 1068, df = 7, *p* < 0.001; Figure 2*b*ii; Table S2; Table S6).

The number of micropyles was found to vary between populations (Anova: *F* = 9.52, *p* < 0.001; Figure 3*a*ii; Table S2; Table S7), as did micropyle width (Anova: *X*^2^ = 20.7, *p* = 0.002; Figure 3*b*ii; Table S2; Table S8).

### Climatic effects on intraspecific egg trait variation

We used a piecewise structural equation model (pSEM) to test whether local climate could explain variation in egg pore traits found across the European latitudinal gradient. We modelled all possible associations between climatic variables and egg traits, as well as all possible associations between egg traits with one another. This initial model included 31 paths, 10 of which were identified as significant after model selection. This analysis revealed that climate was a better predictor of egg traits than egg traits among one another (Figure 4). None of the egg traits were associated with one another (Figure 4). Although all the tested variables improved model fit (reduced the CIC value), maximum temperature and growing days had stronger effects than precipitation or temperature seasonality (Figure 4 and Table S4).

**Figure 4.**
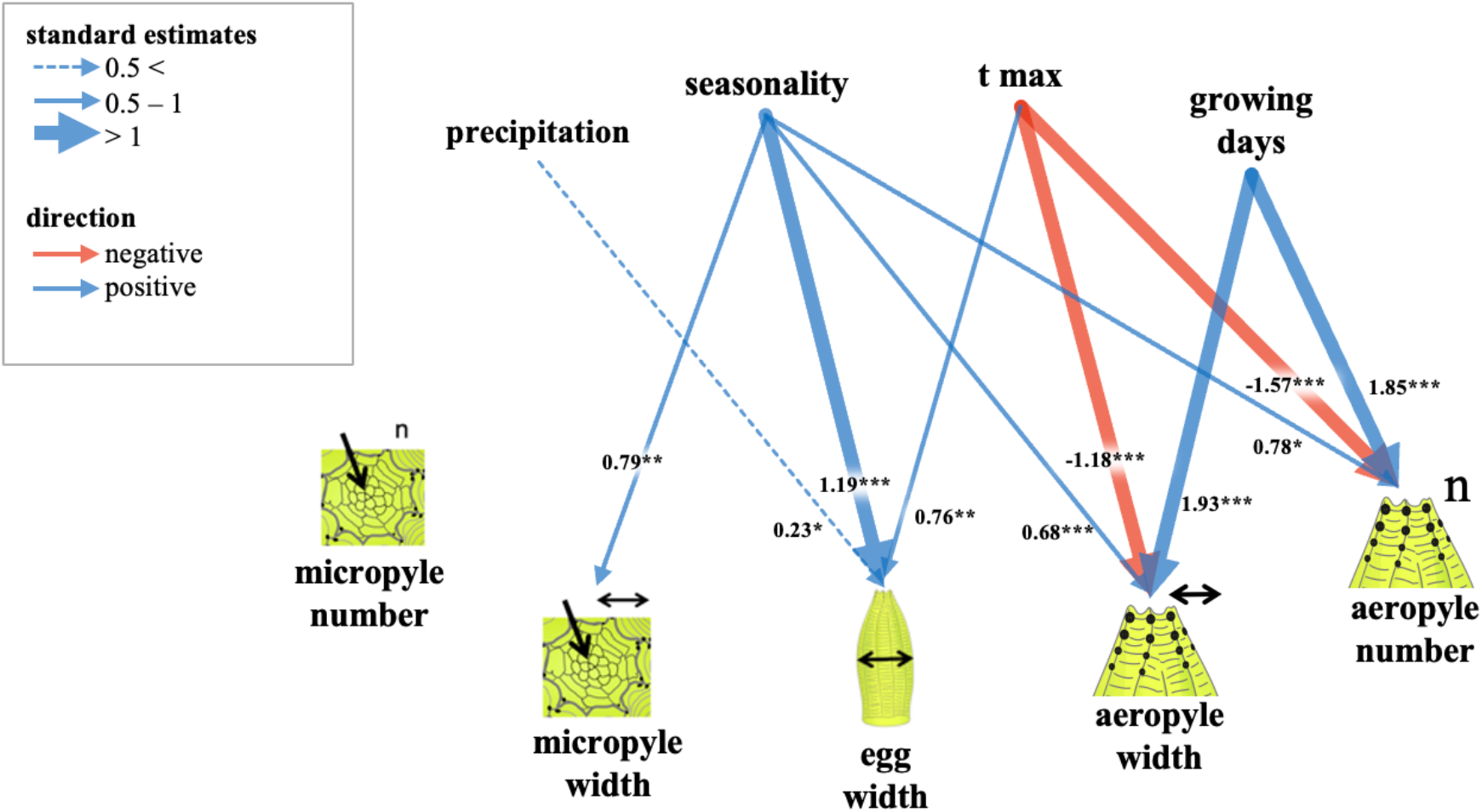
Piecewise structural equation model (p-SEM) indicating significant associations between the climatic variables and the egg traits. The model includes all measured eggs from the seven populations of *Pieris napi* (n = 155 eggs, n = 165 micropyles, n = 4361 aeropyles). Numbers are standard estimates, and asterisks indicate statistical significance (* *p* < 0.05, ** *p* < 0.01, *** *p* < 0.001; Table S4).

### Cover of micropylar pit

For a proportion of the eggs, the micropylar pit was covered by an unknown substance (Figure 3*c*). This covering was found for eggs of all three *Pieris* spp. and for all seven populations of *P. napi*. The cover varied between fully enveloping the micropylar pit, making all features indistinguishable, to a thin or partial layer. Eggs that were imaged less than two hours after egg deposition were covered significantly less often than eggs older than four hours (*glm*: estimate = 1.56, *se* = 0.58, *z* = 2.17, *p* = 0.030; Figure 3*d*).

## Discussion

Our study on egg pore traits (aeropyles and micropyles) in *Pieris* butterflies showed inter- and intra-specific variation in both the number and size of aeropyles and micropyles. Interestingly, *P. napi* populations from divergent climates displayed even greater differentiation than between species, suggesting that climate-associated selection may play a role in shaping these egg pores that are associated with respiration and reproduction. However, given the complexity of potential selective pressures, we emphasise that climate is only one possible driver of this divergence.

An intraspecific comparison of *P. napi* eggs from various populations revealed that climatic variables were significantly associated with both aeropyle and micropyle width, independent of egg size in the intraspecific comparison. Moreover, we observed that *Pieris* eggs often show a secretion covering the micropylar pit within hours of oviposition. This covering may serve to protect against water loss, though its functional role remains speculative. Taken together, our findings suggest that egg pore traits vary within and between *Pieris* species and are potentially responsive to environmental variation. However, determining whether climate causally drives morphological change requires further experimental and genetic work. These egg pores, especially in the context of varying environmental pressures, deserve further attention in studies of insect adaptation to environmental conditions.

### 1. Drivers of interpecific variation of egg pores in *Pieris*

We explored the interspecific morphological variation of egg pores in *Pieris* by comparing aeropyles and micropyles between three closely related species: *P. brassicae, P. rapae*, and *P. napi*. These species co-occur in Europe, all using Brassicaceae host plants, but differ in their oviposition behaviour, thermosensitivity, and microhabitat use, which likely influences egg exposure to microclimates^45,47,63^. Eggs laid in clutches by the gregarious *P. brassicae* were significantly larger and had slightly more but smaller aeropyles than the solitary eggs laid by *P. rapae* and *P. napi*. The latter two species laid eggs of similar size with aeropyles of a similar number and width. This result aligns with broader findings on insect egg morphology, where interspecific variation includes differences in aeropyle shape, number, and spatial distribution^5–7^. Although prior studies confirmed the presence of aeropyles in these species, quantifications were lacking^10,14,64.^ While aeropyles are assumed to evolve under abiotic selection, their precise functional role remains underexplored^5^. Developmental plasticity may also play a role, as aeropyle number and width have been observed to change during embryogenesis^65^. However, developmental plasticity did not play a role in our study as all eggs were imaged between 1 and 3-days-old. Eggs laid in clutches by the gregarious *P. brassicae* were significantly larger and had slightly more but smaller aeropyles than the solitary eggs laid by *P. rapae* and *P. napi*. The latter two species laid eggs of similar size with aeropyles of a similar number and width. This result aligns with broader findings on insect egg morphology, where interspecific variation includes differences in aeropyle shape, number, and spatial distribution^5–7^. Although prior studies confirmed the presence of aeropyles in these species, quantifications were lacking^10,14,64.^ While aeropyles are assumed to evolve under abiotic selection, their precise functional role remains underexplored^5^.

Aeropyles facilitate gas exchange, whereas micropyles mediate sperm entry and may also contribute to environmental adaptation^40^. In contrast to aeropyle patterns, micropyles were more similar between *P. brassicae* (gregarious) and *P. rapae* (solitary), while *P. napi* (solitary) eggs had fewer and smaller micropyles, thus not correlating with egg size or oviposition strategy. Micropyle variation, a common feature across Insecta, is often used for taxonomic classification^5,66,67^. Micropyle number has been linked to differences in precipitation experienced across insects^40^. In Lepidoptera, higher levels of polyandry are associated with more micropyles^9^. In this case, all three *Pieris* species are polyandrous^68,69^, though *P. napi* varies in mating frequency across its range^70^. Our data show that *Pieris* species possess micropyle traits typical of other Papilionoidea^40^, and in this case micropyle number did not vary with climate and may instead reflect greater influence from phylogenetic or reproductive history.

Although all individuals were reared under the same greenhouse conditions, the ecological difference between species in thermosensitivity and oviposition suggests that their eggs evolved experiencing different climate conditions in the field^45,47,63^. For example, *P. napi* is strongly thermosensitive and prefers cool, moist oviposition sites, potentially reducing heat and desiccation stress on its eggs relative to migratory *P. rapae*, whose eggs can be found in more variable habitats^45^. Field-experienced microclimates, especially those at the leaf boundary layer, likely differ substantially by oviposition strategy^49^. These microhabitat differences could influence adaptive egg pore traits. Recent studies find that both egg size and shape are best explained by the environment in which eggs are laid, such as in aquatic environments or within hosts^38^. The findings suggest that ecological context, rather than fixed scaling rules, shapes the evolution of insect eggs^38^. Nevertheless, our design only considers macroclimatic differences, which may also be mitigated by different oviposition strategies. For example, differences in oviposition tissue selection may buffer differences in macroclimate by adjusting microclimates^70^. Future work incorporating field microclimate data would help clarify the relative influence of macroclimatic vs. microclimatic pressures.

Ultimately, interspecific comparisons, especially among only three species, are correlational and constrained by confounding factors such as phylogenetic history, oviposition behaviour, host plant use, and developmental timing^72^. Broader phylogenetic analyses suggest some egg trait variation may be explained by phylogenetic signal rather than adaptive divergence^6,20^. Investigating morphological variation within species, such as rapidly evolving *P. napi*, offers greater potential for disentangling ecological from phylogenetic drivers.

### 2. Drivers of intraspecific variation of egg pores in *P. napi*

*Pieris napi* is particularly suited for examining intraspecific divergence, as its non-migratory, locally adapted populations span a broad latitudinal range^73–76^. In our study, egg pore traits showed even greater divergence among *P. napi* populations than between *Pieris* species, suggesting rapid trait evolution. Notably, the extreme populations at the leading and trailing edges of the population range, the northern Abisko and southern Costa Brava, exhibited the most divergent egg pore traits, while mid-range populations were more similar to one another. Egg pore differences among *P. napi* populations persisted after multiple generations in greenhouse conditions, indicating a heritable basis. This persistent difference across generation makes phenotypic plasticity an unlikely sole explanation. Nonetheless, plasticity cannot be excluded entirely. Edge populations, both northern and southern, often show greater plasticity than central populations^77^, raising the possibility that Abisko and Costa Brava eggs exhibit more plastic responses to compensate for being reared under suboptimal conditions. Divergence at the range edges could therefore reflect either adaptive divergence or evolved differences in plasticity depending on the bioclimatic context from which the species was collected^77^.

Genetic drift is a potential explanation for divergence in geographically isolated populations, particularly in species like *P. napi* with limited dispersal. Under a model of isolation-by-distance, we would expect genetic and phenotypic differences to increase gradually with geographic distance due to restricted gene flow among distant populations^78^. Similarly, postglacial recolonisation from southern refugia after the last glacial maximum typically results in gradual, clinal patterns of genetic variation along latitudinal gradients, reflecting sequential founder effects during northward expansion^78^. However, our results reveal a non-linear, polynomial pattern of egg pore trait divergence, where both northernmost (Abisko) and southernmost (Costa Brava) populations show the most extreme phenotypes, while central populations are more similar to one another (Figure 5). This pattern is inconsistent with a simple isolation-by-distance model or unidirectional recolonisation, both of which would predict more gradual change ^78,79^. Instead, the observed pattern suggests that local adaptation or selection-driven divergence, possibly interacting with evolved plasticity, is a more likely driver of trait variation at the range edges. Recent genomic analyses provide support for climate-associated selection, identifying summer temperature and precipitation as key drivers of genetic differentiation in *P. napi*, with Iberian populations (like Costa Brava) and the subspecies *P. napi* adalwinda (like Abisko) as genetic outliers^79^. The same populations show the most pronounced divergence in egg pore morphology in our study. While this strengthens the case for climate-mediated selection, it remains correlative. Experimental approaches, such as testing trait performance under variable climates or associating trait variation with climate-linked loci, are needed to establish causality and adaptive evolution of egg pore traits.

**Figure 5.**
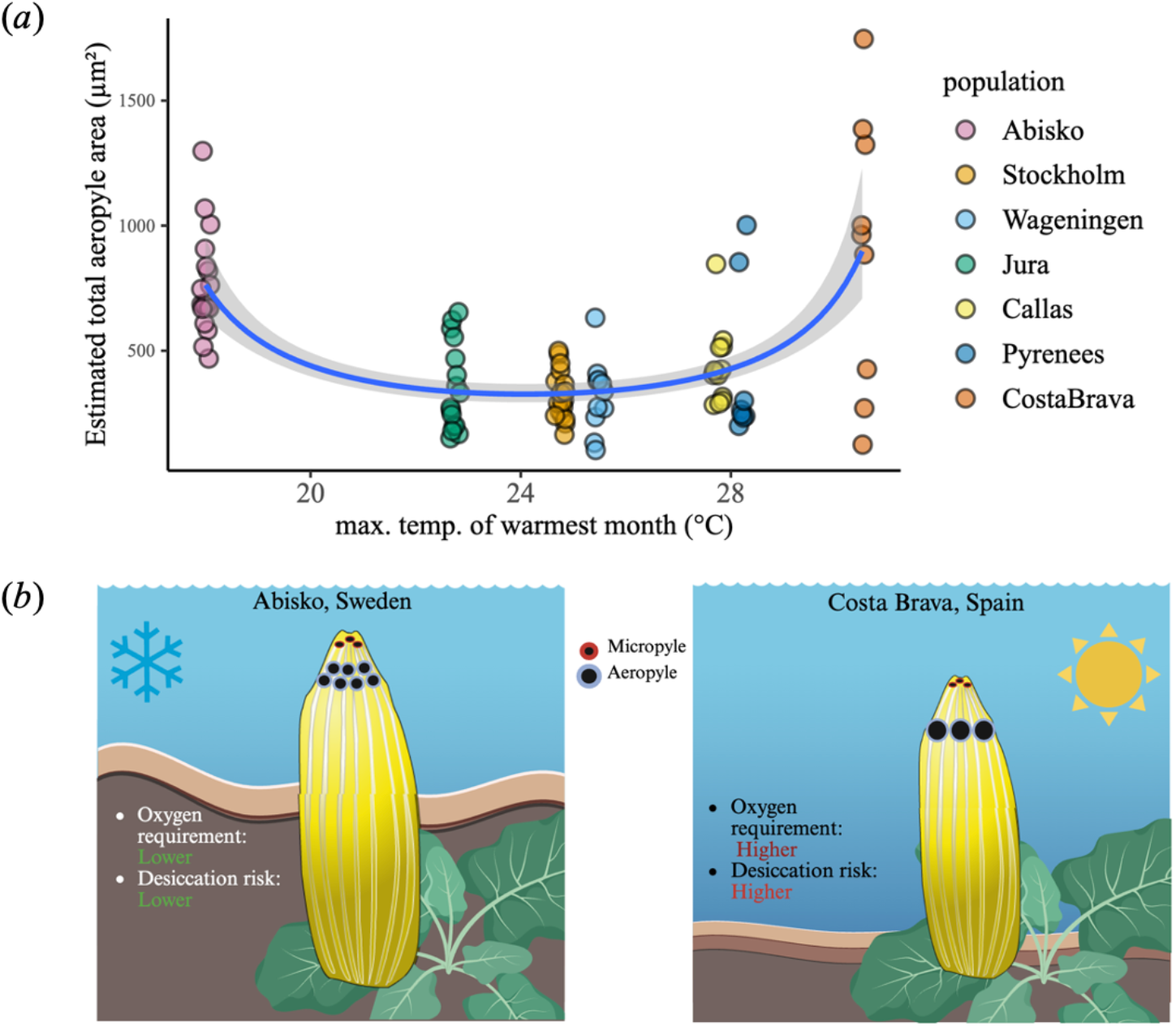
Divergent egg pore morphology in *Pieris napi* across climatic extremes reflects a trade-off between aeropyle number and size. (*a*) Aeropyle area (width × number) plotted against the maximum temperature of the warmest month shows a non-linear divergence, with the greatest area in northern (Abisko) and southern (Costa Brava) populations, which occupy the coldest and hottest parts of the species range, respectively. The trendline was plotted using a glm with the formula y∼poly(x,2) and Gamma distribution. (*b*) Schematic summary: in Abisko, increased aeropyle area results from more, smaller aeropyles and is paired with larger eggs. In Costa Brava, larger but fewer aeropyles likely meet higher respiratory demands while minimising water loss; eggs are laid on plants near water to buffer against heat and dryness.

Aeropyle number and width varied markedly among *P. napi* populations and correlated with thermal extreme, even more than between the species. Costa Brava eggs, exposed to the hottest climates, had the widest aeropyles, potentially to support increased oxygen demand under elevated metabolic rates, similar to patterns observed in the seed beetle *Callosobruchus maculatus*, where higher larval metabolism correlates with larger aeropyles^22,80^. In contrast, Abisko eggs had the most aeropyles, which may reflect relaxed selection against water loss in its cool, humid environment. These patterns suggest that thermal conditions drive divergence in pore morphology through dual constraints: oxygen demand and desiccation risk. Total gas exchange capacity appears maintained across populations via different trait combinations, indicating functional trade-offs: many small aeropyles (Abisko) versus fewer large ones (Costa Brava).

Micropyle traits also varied among populations, with Abisko eggs having the most and largest micropyles. While intraspecific micropyle variation is less common than interspecific divergence^40,81^, *P. napi* populations displayed divergence on par with that among closely related *Pieris* species, suggesting greater evolutionary flexibility than previously appreciated. However, only micropyle width, not number, was significantly associated with a climatic variable, seasonality. Overall, micropyles exhibited less climate-linked divergence than aeropyles, possibly due to stronger functional constraints or differing selective pressures related to reproduction.

Egg size also varied, with Abisko eggs significantly larger than those from other populations, consistent with Bergmann’s rule and patterns seen in other Lepidoptera^28,82^. This contradicts the “canteen hypothesis,” which predicts larger eggs in dry environments to buffer against desiccation, a hypothesis developed in *Drosophila* (Diptera) and so far, largely unsupported in Lepidoptera^26^. Instead, colder temperatures likely favour larger eggs for developmental benefits; for example, butterflies experimentally shifted to colder conditions have been shown to lay larger eggs, improving offspring survival^82^. Despite size differences, aeropyle area did not scale with egg size. Instead, aeropyle traits were more strongly predicted by climatic variables, particularly maximum temperature and growing degree days, factors known to drive divergence traits such as larval respiration rates in other Lepidoptera^29^. This decoupling suggests that eggs may compensate for environmental demands not by scaling, but through structural or physiological adaptations such as altered shell permeability or enhanced gas exchange via the vitelline membrane or serosa^23,83^.

The absence of extreme egg traits in *P. napi* mid-range populations aligns with findings that populations near the centre of their thermal niche often experience more stable environments and exhibit reduced physiological divergence^77^. Still, while climate appears to be a major driver of variation, other ecological variables, such as host plant traits, predation pressure, or microhabitat variation, could also shape egg trait evolution^84^. Moreover, protective egg structures like the serosa, which helps prevent desiccation^84^, may interact with changes in egg pore morphology in ways not yet understood. Demonstrating adaptive significance will ultimately require fitness-based experiments across environmental gradients.

### 3. Micropyle Coating: A Potential Strategy Against Abiotic and Biotic Stress

We found that several hours after oviposition, micropyles were no longer visible on many eggs. By comparing freshly laid eggs to those at least four hours old, we observed that this covering was dynamic and changed over time. The covering may have been secreted several hours after oviposition, or it was already deposited as a transparent substance at the time of oviposition and hardened over time, becoming visible. In contrast to most insect eggs, the inner surface of *P. brassicae* eggs is lined with a layer of unsaturated oil over the region surrounding the micropyle^14^. This oil layer flows into the area of the micropyles within four hours of oviposition and is later reabsorbed before eclosion^14^. Experimental studies show that *P. brassicae* eggs are only susceptible to water-soluble poisons during this brief window, suggesting that the oily coating plays a protective role that the eggshell alone cannot provide^14^. It is likely that the gradual micropyle covering we observed corresponds to the previously described oily layer.

In addition to blocking toxin uptake, non-volatile oily materials may interfere with respiration by obstructing the air spaces in the shell^14^. Our findings, together with prior work, suggest that micropyle traits vary with climate, potentially influencing how embryos interact with environmental conditions^40^. If micropyles indeed mediate climatic effects, this temporary coating may serve to shield embryos from environmental variation. Analogous systems support this interpretation: hydrogel-based egg coatings protect against drought^44^ and heat stress^8^, while oil layers may reduce water loss through micropyles that would otherwise remain open. Notably, insect eggs can exchange gases through micropyles^41,42^, and coatings could help regulate this process. Similar protective layers have been described in aquatic eggs of fish^85^ and insects^43^, and a waxy chorionic layer in terrestrial Hemiptera also serves a waterproofing function^11,86^. These parallels suggest convergent evolution of protective coatings in diverse taxa. Whether the oily layer in *Pieris* functions analogously remains an open question deserving further investigation.

Beyond preventing water loss, egg coatings may also protect against solar radiation and associated temperature or UV stress^87^. Intriguingly, they may also offer defence against biotic threats. Egg parasitoid wasps locate hosts via chemical cues^88^, and coatings might mask or disrupt these cues or form a physical barrier to oviposition^44^. Whether such coatings influence the risk of egg parasitism in *Pieris* remains unexplored.

While speculative, these hypotheses highlight a potentially underappreciated layer of egg adaptation. Future work should investigate whether micropylar coatings vary across climates and whether they contribute to egg survival under different abiotic or biotic stresses.

## Conclusion

Our study reveals that egg pores in *Pieris* butterflies are highly variable traits shaped by a complex interplay of ecological, behavioural, and climatic factors. Across both inter- and intraspecific comparisons, we found variation in aeropyle traits, and to a lesser extent, in micropyles, with particularly pronounced divergence among *P. napi* populations from climatically extreme regions.

These differences persisted across multiple generations in greenhouse conditions, indicating a heritable component. Correlations with climate variables such as temperature and growing season length support the hypothesis of local adaptation. Recent phylogeographic genomic analyses identifying climate-associated population divergence in *P. napi* genomes strengthen the case that climatic selection may play a role^79^. The observation that Abisko and Costa Brava populations differ most both in climate and in egg pore morphology suggests that egg traits may be part of a broader adaptive response to local environments. However, genetic drift, developmental plasticity, or other ecological pressures may also contribute to the observed patterns.

Our results also highlight a potentially under-recognised layer of egg adaptation: the coating that forms over the micropylar pit. This secretion may help buffer embryos against desiccation, temperature stress, or parasitism, particularly in climates where such pressures are acute. The potential interaction between micropyle width, environmental exposure, and post-oviposition sealing represents a promising avenue for future research.

Together, these findings emphasise that insect egg traits are not merely static or taxonomic features but may be dynamic, responsive elements of the reproductive phenotype. Understanding their evolution will require integrative approaches that link morphology, physiology, microhabitat selection, and climatic context. As climates continue to shift, such traits may play an increasingly important role in shaping reproductive success and species persistence.

## References

1. Gibbs, M. & Dyck, H. Van. Reproductive plasticity, oviposition site selection, and maternal effects in fragmented landscapes. Behavioral Ecology and Sociobiology (2009). 10.1007/s00265-009-0849-8

2. Potter, K., Goggy Davidowitz & H. Arthur Woods. Insect eggs protected from high temperatures by limited homeothermy of plant leaves. (2009). 10.1242/jeb.033365

3. Schäpers, A., Nylin, S., Carlsson, M. A. & Niklas Janz. Specialist and generalist oviposition strategies in butterflies: maternal care or precocious young? Oecologia 1, 335– 343 (2016). 10.1007/s00442-015-3376-5

4. Tourneur, J.-C., Cole, C., Vickruck, J., Dupont, S. & Meunier, J. Pre-and post-oviposition behavioural strategies to protect eggs against extreme winter cold in an insect with maternal care. Journal of Experimental Biology. (2022). 10.24072/pcjournal.104

5. Hinton, H. E. Biology of Insect Eggs. Biology of Insect Eggs 1, 1–1111 (2013). 10.1016/c2013-1-15220-3

6. Regier, J. C., Paukstadt, U., Paukstadt, L. H., Mitter, C. & Peigler, R. S. Phylogenetics of Eggshell Morphogenesis in Antheraea (Lepidoptera: Saturniidae): Unique Origin and Repeated Reduction of the Aeropyle Crown. Syst. Biol 54, 254–267 (2005). 10.1080/10635150590923281

7. E. S. Omelina, E. M. B. Main Types of Respiratory System Structure of Eggshells in Insects and Genes Participating in Their Development. Biology Bulletin Reviews. (2013). 10.1134/s2079086413010076

8. Hilker, M., Salem, H. & Fatouros, N. E. Adaptive Plasticity of Insect Eggs in Response to Environmental Challenges. Annual Review of Entomology. 252, 54 (2023). 10.1146/annurev-ento-120120-100746

9. Iossa, G., Gage, M. J. G. & Eady, P. E. Micropyle number is associated with elevated female promiscuity in Lepidoptera. Biol Lett 12, (2016). 10.1098/rsbl.2016.0782

10. Telfer, W. H. Egg formation in Lepidoptera. Telfer Journal of Insect Science. 9, (2009). 10.1098/rsbl.2016.0782

11. Beament, J. W. L. The formation and structure of the micropylar complex in the egg-shell of Rhodnius prolixus stahl. (Heteroptera: Reduviidae). Journal of Experimental Biology. (1947). 10.1242/jeb.23.3-4.213

12. Kubrakiewicz, J., Jȩdrzejowska, I., Szymańska, B. & Biliński, S. M. Micropyle in neuropterid insects. Structure and late stages of morphogenesis. Arthropod Struct Dev 34, 179–188 (2005). 10.1016/j.asd.2005.02.001

13. Campbell, B. E. et al. Complications with Controlling Insect Eggs. Insecticides Resistance (2016). 10.5772/61848

14. Beament, J. W. L. & Lal, R. Penetration through the Egg-shell of Pieris brassicae (L.). Bull Entomol Res 48, 109–125 (1957). 10.1017/s0007485300054134

15. Gautam, S. G., Opit, G. P., Margosan, D., Tebbets, J. S. & Walse, A. S. Egg Morphology of Key Stored-Product Insect Pests of the United States. Ann. Entomol. Soc. Am 107, 1–10 (2014). 10.1603/an13103

16. Kaleka, A. S., Jallundhara, S. & Kapoor, Y. Scanning Electron Microscope Studies on Ornamentation of Egg Chorion of Capissa vagesa (Moore, 1859) (Erebidae) and Trabala vishnou (Lefèbvre, 1827) (Lasiocampidae) (Ditrysia: Lepidoptera) from India. J Entomol Res Soc 25, 337–349 (2023). 10.51963/jers.2023.92

17. Koss, R. Morphology and Taxononiic Use of Ephemeroptera Eggs. Annals of the Entomological Society of America. (1968). 10.1093/aesa/61.3.696

18. Zeh, D. W., Zeh, J. A. & Smith, R. L. Ovipositors, amnions and eggshell architecture in the diversification of terrestrial arthropods. Quarterly Review of Biology 64, 147–168 (1989). 10.1086/416238

19. Hinton, H. E. Function of shell structures of pig louse and how egg maintains a low equilibrium temperature in direct sunlight. J Insect Physiol 23, 785–800 (1977). 10.1016/0022-1910(77)90001-4

20. Boisseau, R. P. & Woods Correspondence, H. A. Resource allocation strategies and mechanical constraints drive the diversification of stick and leaf insect eggs. Current Biology 34, 2880-2892.e7 (2024). 10.1016/j.cub.2024.05.042

21. Donoughe, S. Insect egg morphology: evolution, development, and ecology. Curr Opin Insect Sci 50, 100868 (2022). 10.1111/j.1365-3032.1994.tb01070.x

22. Daniel, S. & Smith, R. H. Functional anatomy of the egg pore in Callosobruchus rnaculatus: a trade-off between gas-exchange and protective functions? Physiological Entomology 19, 30–38 (1994). 10.1111/j.1365-3032.1994.tb01070.x

23. Woods, H. A. Water loss and gas exchange by eggs of Manduca sexta: Trading off costs and benefits. J Insect Physiol 56, 480–487 (2010). 10.1016/j.jinsphys.2009.05.020

24. Woods, H. A. & Bonnecaze, R. T. Insect eggs at a transition between diffusion and reaction limitation: Temperature, oxygen, and water. J Theor Biol 243, 483–492 (2006). 10.1016/j.jtbi.2006.07.008

25. Woods, H.A.* and Ryan I. Hill†. Temperature-dependent oxygen limitation in insect eggs. Journal of Experimental Biology. (2004). 10.1242/jeb.00991

26. Potter, K. A. & Woods, H. A. No evidence for the evolution of thermal or desiccation tolerance of eggs among populations of Manduca sexta. Funct Ecol 26, 112–122 (2012). 10.1111/j.1365-2435.2011.01912.x

27. Parry, D. Macrogeographic clines in fecundity, reproductive allocation, and offspring size of the forest tent caterpillar Malacosoma disstria. Ecological Entomology. (2001). 10.1046/j.1365-2311.2001.00319.x

28. Garcia-Barros, E. Body size, egg size, and their interspecific relationships with ecological and life history traits in butterflies (Lepidoptera: Papilionoidea, Hesperioidea). (1999). 10.1111/j.1095-8312.2000.tb00210.x

29. Ayres, M. P. & Mark Scriber, J. Local adaptation to regional climates in Papilio Canadensis (lepidoptera: papilionidae). Ecol Monogr 64, 465–482 (1994). 10.2307/2937146

30. Horne, C. R., Hirst, A. G., Atkinson, D. & Andrew Hirst, C. G. Insect temperature-body size trends common to laboratory, latitudinal and seasonal gradients are not found across altitudes. Functional Ecology. (2017) 10.1111/1365-2435.13031

31. Meiri, S. & Dayan, T. On the validity of Bergmann’s rule. J Biogeogr 30, 331–351 (2003). 10.1046/j.1365-2699.2003.00837.x

32. Shelomi, M. Where Are We Now? Bergmann’s Rule Sensu Lato in Insects. The American Naturalist 180, (2012). 10.1086/667595

33. Alcantara, M.J.M., Fontanilla, A.M., Ashton, L.A., Burwell, C.J., Cao, M., Han, H., Huang, H., Kitching, R.L., Reshchikov, A., Shen, X. and Tang, Y. Bugs and Bergmann’s rule: a cross-taxon large-scale study reveals idiosyncratic altitudinal and latitudinal body size patterns for different insect taxa. Entomol. Gen. (2024). 10.1127/entomologia/2024/2246

34. Pimentel, C., Ferreira, C. & Nilsson, J.-Å. Latitudinal gradients and the shaping of life-history traits in a gregarious caterpillar. Biological Journal of the Linnean Society. (2010). 10.1111/j.1095-8312.2010.01413.x

35. Gibbs, A. & Matzkin, L. Evolution of water-balance in Drosophilla. Journal of Experimental Biology. (2001). 10.1242/jeb.204.13.2331

36. Clark, B. R. & Faeth, S. H. The evolution of egg clustering in butterflies: A test of the egg desiccation hypothesis. Evolutionary Ecology. (1998). 10.1023/A:1006504725592

38. Church, S. H., Donoughe, S., de Medeiros, B. A. S. & Extavour, C. G. Insect egg size and shape evolve with ecology but not developmental rate. Nature 571, 58–62 (2019). 10.1038/s41586-019-1302-4

39. Mongue, A. J., Megan E. Hansen, Liuqi Gu, Clyde E. Sorenson & James R. Walters. Nonfertilizing sperm in Lepidoptera show little evidence for recurrent positive selection. Molecular Ecology. (2018). 10.1111/mec.15096

40. Iossa, G.The ecological function of insect egg micropyles. Funct Ecol 36, 1113–1123 (2022). 10.1111/1365-2435.14023

41. Capinera, J. L., Naranjo, S. E. & Packard, M. J. Vapor Density and Water Loss from Eggs of the Range Caterpillar, Hemileuca oliviae. (1981). 10.1093/ee/10.1.97

42. Tuft, P. H. The structure of the insect egg-shell in relation to the respiration of the embryo. Journal of Experimental Biology. 26, (1950). 10.1242/jeb.26.4.327

43. Barbier, R. & Chauvin, G. The aquatic egg of Nymphula nymphaeata (Lepidoptera: Pyralidae) - On the fine structure of the egg shell. Cell Tissue Res 149, 473–479 (1974). 10.1007/bf00223026

44. Li, T. H., Wang, X., Desneux, N., Wang, S. & Zang, L. S. Egg coverings in insects: ecological adaptation to abiotic and biotic selective pressures. Biological Reviews 100, 99–112 (2025). 10.1111/brv.13130

45. Vives-Ingla, M., Sala-Garcia, J., Stefanescu, C. & Peñuelas, J. Varying thermal exposure, host-plant traits and oviposition behaviour across vegetation ecotones. BioRxiv. (2020) 10.1101/2020.02.11.944439

46. Friberg, M. & Wiklund, C. Host preference variation cannot explain microhabitat differentiation among sympatric Pieris napi and Pieris rapae butterflies. Ecol Entomol 44, 571–576 (2019). 10.1111/een.12728

47. Tolman, T. & Lewington, R. Butterflies of Britain and Europe. Collins Field Guide. (2009).

48. Barton, B. T. & Schmitz, O. J. Experimental warming transforms multiple predator effects in a grassland food web. Ecol Lett 12, 1317–1325 (2009). 10.1111/j.1461-0248.2009.01386.x

49. Pincebourde, S. & Woods, H. A. Climate uncertainty on leaf surfaces: the biophysics of leaf microclimates and their consequences for leaf-dwelling organisms. Functional Ecology. (2012). 10.1111/j.1365-2435.2012.02013.x

50. Schindelin, J. et al. Fiji: An open-source platform for biological-image analysis. Nat Methods 9, 676–682 (2012). 10.1038/nmeth.2019

51. Fick, S. E. & Hijmans, R. J. WorldClim 2: new 1-km spatial resolution climate surfaces for global land areas. Int. J. Climatol (2017) doi:10.1002/joc.5086.

52. Zaman, K., Hubert, M. K. & Schoville, S. D. Testing the role of ecological selection on colour pattern variation in the butterfly Parnassius clodius. Mol Ecol 28, 5086–5102 (2019). 10.1111/mec.15279

53. Beerli, N., Bärtschi, F., Ballesteros-Mejia, L., Kitching, I. J. & Beck, J. How has the environment shaped geographical patterns of insect body sizes? A test of hypotheses using sphingid moths. J Biogeogr 46, 1687–1698 (2019). 10.1111/jbi.13583

54. Cayton, H. L., Haddad, N. M., Gross, K., Diamond, S. E. & Ries, L. Do growing degree days predict phenology across butterfly species? Ecology 96, 1473–1479 (2015). 10.1890/15-0131.1

55. Fischer, Günther, Freddy O. Nachtergaele, H. Van Velthuizen, F. Chiozza, G. Francheschini, M. Henry, D. Muchoney, and S. Tramberend. Global agro-ecological zones (gaez v4)-model documentation. (2021). 10.4060/cb4744en

56. Robert J. Hijmans. Package: dismo. (2024). https://rspatial.org/raster/sdm/.

57. Harris, I., Osborn, T. J., Jones, P. & Lister, D. Version 4 of the CRU TS monthly high-resolution gridded multivariate climate dataset. (2020) 10.1038/s41597-020-0453-3

58. R Core Team. R: A Language and Environment for Statistical Computing. R Foundation for Statistical Computing, (2020). https://www.r-project.org

59. Bates, D., Mächler, M., Bolker, B. M. & Walker, S. C. Fitting Linear Mixed-Effects Models Using lme4. J Stat Softw 67, 1–48 (2015). 10.18637/jss.v067.i01

60. Hartig, F. Residual Diagnostics for Hierarchical (Multi-Level / Mixed) Regression Models [R package DHARMa version 0.4.7]. CRAN: Contributed Packages (2024).

61. Lenth, R. V. emmeans: Estimated Marginal Means, aka Least-Squares Means. CRAN: Contributed Packages (2017). 10.32614/cran.package.emmeans

62. Lefcheck, J. S. piecewiseSEM: Piecewise structural equation modelling in r for ecology, evolution, and systematics. Methods Ecol Evol 7, 573–579 (2016). 10.1111/2041-210x.12512

63. Friberg, M. & Wiklund, C. Host preference variation cannot explain microhabitat differentiation among sympatric Pieris napi and Pieris rapae butterflies. Ecological Entomology. (2019). 10.1111/een.12728

64. Feltwell, J. Large White Butterfly. Large White Butterfly (1981). 10.1007/978-94-009-8638-1

65. Renthlei, C. Z., Raghuvarman, A., Kharbuli, B. & Dey, S. Progressive chorion morphology during egg development in Samia ricini (Donovan). Microsc Res Tech 73, 234–239 (2010). 10.1002/jemt.20781

66. R. H. Cobben. Evolutionary Trends in Heteroptera. Part I. Eggs, Architecture of the Shell, Gross Embryology and Eclosion. Wageningen: Centre for Agricultural Publishing and Documentation (1968). 10.2307/2406799

67. Trougakos, I. P. & Margaritis, L. H. Novel Morphological and Physiological Aspects of Insect Eggs. Chemoecology of Insect Eggs and Egg Deposition 2–36 (2003). 10.1002/9780470760253.ch1

68. David, W. A. L. & Gardiner, B. O. C. The mating behaviour of Pieris brassicae (L.) in a laboratory culture. Bull Entomol Res 52, 263–280 (1961). 10.1017/s0007485300055401

69. Bissoondath, C. J. & Wiklund, C. Effect of male mating history and body size on ejaculate size and quality in two polyandrous butterflies, Pieris napi and Pieris rapae (Lepidoptera : Pieridae). Funct Ecol 10, 457–464 (1996). 10.2307/2389938

70. Bergström, J., Wiklund, C. & Kaitala, A. Natural variation in female mating frequency in a polyandrous butterfly: effects of size and age. Anim Behav 64, 49–54 (2002). 10.1006/anbe.2002.3032

71. Melero, Y., Evans, L.C., Kuussaari, M., Schmucki, R., Stefanescu, C., Roy, D.B. and Oliver, T.H., 2022. Local adaptation to climate anomalies relates to species phylogeny. Communications Biology (2022). 10.1038/s42003-022-03088-3

72. Okamura, Y. et al. Microevolution of Pieris butterfly genes involved in host plant adaptation along a host plant community cline. Mol Ecol 31, 3083–3097 (2022). 10.1111/mec.16447

73. Keehnen, N. L. P., Hill, J., Nylin, S. & Wheat, C. W. Microevolutionary selection dynamics acting on immune genes of the green-veined white butterfly, Pieris napi. Mol Ecol 27, 2807–2822 (2018). 10.1111/mec.14722

74. Keehnen, N.L., Fors, L., Järver, P., Spetz, A.L., Nylin, S., Theopold, U. and Wheat, C.W., 2019. Geographic variation in hemocyte diversity and phagocytic propensity shows a diffuse genomic signature in the green veined white butterfly. bioRxiv. (2019). 10.1101/790782

75. Neethiraj, R., Pruisscher, P., Pruisscher Keehnen, N., Woronik, A., Gotthard, K., Nylin, S. and Wheat, C. A dark melanic morph of Pieris napi shares its origins with other dark morphs of Lepidoptera. (2019).

76. Evans, L.C., Melero, Y., Schmucki, R., Boersch‐Supan, P.H., Brotons, L., Fontaine, C., Jiguet, F., Kuussaari, M., Massimino, D., Robinson, R.A. and Roy, D.B. Bioclimatic context of species’ populations determines community stability. Global Ecology and Biogeography 31, 1542–1555 (2022). 10.1111/geb.13527

77. Vandewoestijne, S. & Dyck, H. Van. Population Genetic Differences along a Latitudinal Cline between Original and Recently Colonized Habitat in a Butterfly. Plos one. (2010). 10.1371/journal.pone.0013810

78. Sala-Garcia, J. et al. Phylogeography and diversification of the Pieris napi species group in the Western Palaearctic. bioRxiv (2025). 10.1101/2025.01.31.634921

79. Gillooly, J. F., Charnov, E. L., West, G. B., Savage, V. M. & Brown, J. H. Effects of size and temperature on developmental time. Nature 417, 70–73 (2002). 10.1101/2025.01.31.634921

80. Matesco, V. C. et al. External egg structure of the Pentatomidae (Hemiptera: Heteroptera) and the search for characters with phylogenetic importance. Zootaxa 3768, 351–385 (2014). 10.11646/zootaxa.3768.3.5

81. Fischer, K., Brakefield, P. M. & Zwaan, B. J. Plasticity in butterfly egg size: Why larger offspring at lower temperatures? Ecology 84, 3138–3147 (2003). 10.1890/02-0733

82. Zrubek, B. & Woods, H. A. Insect eggs exert rapid control over an oxygen-water tradeoff. Proceedings of the Royal Society B: Biological Sciences 273, 831–834 (2006). 10.1098/rspb.2005.3374

83. Harvey, J. A., Heinen, R., Gols, R. & Thakur, M. P. Climate change-mediated temperature extremes and insects: From outbreaks to breakdowns. Glob Chang Biol 26, 6685–6701 (2020). 10.1111/gcb.15377

84. Kunz, Y. Developmental Biology of Teleost Fishes. (2004). 10.1007/978-1-4020-2997-4

85. J. W. L. Beament. The waterproofing Process in Eggs of Rhodnius prolixus Stáhl. Proceedings of the Royal Society of London. Series B-Biological Sciences. (1946). 10.1007/978-1-4020-2997-4

86. Hinton, H. E. Function of shell structures of pig louse and how egg maintains a low equilibrium temperature in direct sunlight. J Insect Physiol 23, 785–800 (1977). 10.1016/0022-1910(77)90001-4

87. Gautschi, W. Orthogonal polynomials: Applications and computation Article in Acta Numerica. (1996). 10.1017/S0962492900002622

88. Alejandro Gonzalez-Voyer and Achaz von Hardenberg. An Introduction to Phylogenetic Path Analysis. (2014). 10.1007/978-3-662-43550-2_8

