## Supplementary materials for "Butterfly egg plasticity in physiologically and reproductive relevant traits: adaptation to climate?"

### Content


1. Egg width as a proxy for volume
   - Figure S1. Egg width and length as proxy of egg volume in *Pieris napi*.
   - Table S1. Comparing photograph-measured egg width to cyro-Scanning Electron Microscopy (SEM) measured egg width per population ($\bar{x}$ ± SD (n)).
2. Summarised egg-measurements per *Pieris* rearing
   - Table S2. Measurements for egg characteristics ($\bar{x}$ ± SD (n)) for the three *Pieris* spp. (location = Wageningen) and the seven *P. napi* populations.
   - Figure S2. Comparison of egg width interspecifically between Pieris spp. and intraspecifically between P. napi populations.
3. Details of the seven *P. napi* populations
   - Table S3. Summarized conditions for each of the European populations
4. Correlation plots between egg traits
   - Figure S3. Relations between climatic variables and egg pore traits.
5. Piecewise Structural Equation Model (pSEM) selection
   - Figure S4. Phase one of model selection, model reduction.
   - Figure S5. Phase two of model selection, testing alternative causal hypotheses.
   - Table S4. Standard estimates, standard errors, degrees of freedom, critical values and p-values for the final model.
6. Pairwise comparisons for population egg traits
   - Table S5. Aeropyle number compared between the seven *Pieris napi* populations.
   - Table S6. Aeropyle width compared between the seven *Pieris napi* populations.
   - Table S7. Micropyle number compared between the seven *Pieris napi* populations.
   - Table S8. Micropyle width compared between the seven *Pieris napi* populations.
   - Table S9. Egg width compared between the seven *Pieris napi* populations.


#### 1 Egg width as proxy for volume

To validate the use of egg width as a proxy for egg size, we measured the width and length of 72 eggs from four *P. napi* populations (Abisko, Wageningen, Jura, and Costa Brava). Butterflies were reared as described in ‘Material and Methods’ section ‘insects and plants’, and eggs were collected in February 2025. Eggs were photographed using a Leica DM i1 microscope at five times magnification.

The average egg width per population was very similar to the average egg width measured from the cyro-Scanning Electron Microscope images, indicating that egg size remains relatively constant between generations (Table S1).

Egg volume was estimated by using the formula for an ellipsoid; *Volume* = *(π/6)*egg_length*egg_width^2*[28]. We found egg width to be a good proxy for estimated egg volume (*lm()* not correcting for population; Adjusted R-squared = 0.91; see Figure S1).

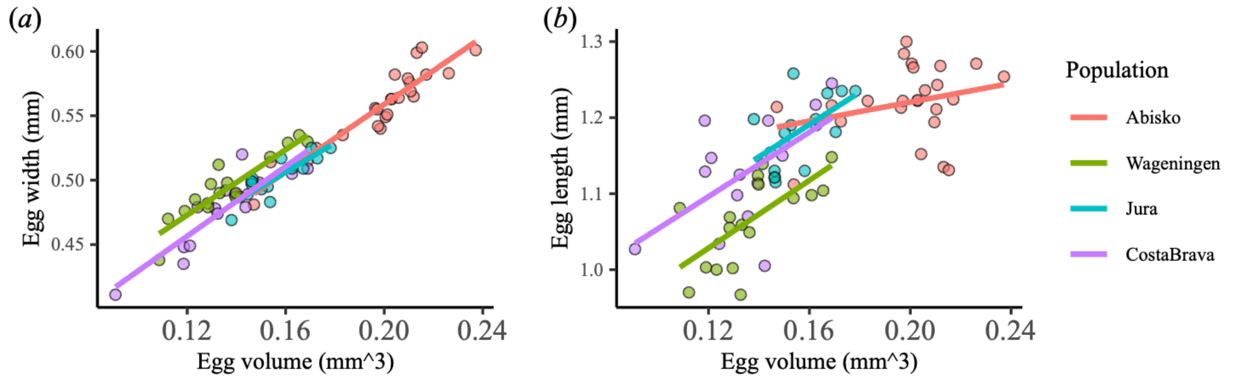


**Figure S1. Egg width (*a*) and length (*b*) as proxy of egg volume in *Pieris napi*.** Egg width showed a good fit to the egg volume of 72 eggs from four *P. napi* populations. Egg volume was estimated by the formula for an ellipsoid: *Volume = (π/6)*length*width^2* [28].

**Table S1. Comparing photograph-measured egg width to cyro-Scanning Electron Microscopy (SEM) measured egg width per population ($\bar{x}$ ± SD (n)).** Averages per population were similar, indicating there is only limited plasticity in egg size between *P. napi* generations. All photographs were taken in 2025. The SEM-measured eggs were either taken in 2023 (Wageningen, Jura and Costa Brava populations) or in 2024 (Abisko population).

| **Population** | **Egg length (µm) – photograph-measured** | **Egg width (µm) – photograph-measured** | **Egg width (µm) – SEM-measured** |
|---|---|---|---|
| Abisko | 1220 ± 49 µm (24) | 558 ± 39.8 µm (14) | 531 ± 15 (25) |
| Wageningen | 1150 ± 67 µm (17) | 477 ± 31.1 µm (14) | 463 ± 24 µm (25) |
| Jura | 1066 ± 57 µm (18) | 495 ± 25.0 µm (11) | 489 ± 29 µm (21) |
| Costa Brava | 1185 ± 48 µm (13) | 503 ± 16.4 µm (11) | 488 ± 25 µm (19) |

### 2 Summarised egg-measurements per *Pieris* rearing

**Table S2. Measurements for egg characteristics ($\bar{x}$ ± SD (n)) for the three *Pieris* spp. (location = Wageningen) and the seven *P. napi* populations**. Numbers in brackets are sample sizes (eggs). For *Pieris brassicae*, one sample consists of one egg clutch, measured as five random eggs from the clutch.

| **A) *Pieris* Species** | **Egg width (µm)** | **Aeropyle number** | **Aeropyle width** | **Micropyle number** | **Average micropyle width** |
|---|---|---|---|---|---|
| *P. brassicae* | 575 ± 22 µm (14) | 47 ± 8.3 (14) | 2.16 ± 0.31 µm (14) | 3.4 ± 0.65 (14) | 1.35 ± 0.18 µm2 (14) |
| *P. rapae* | 462 ± 26 µm (17) | 42 ± 9.2 (14) | 3.24 ± 0.58 µm (17) | 3.5 ± 0.76 (8) | 1.21 ± 0.38 µm2 (8) |
| *P. napi* (Wag.) | 463 ± 31 µm (25) | 43 ± 11.1 (11) | 2.73 ± 0.47 µm (25) | 2.2 ± 0.86 (15) | 0.87 ± 0.22 µm2 (15) |
| **B) *P. napi* Population** | | | | | |
| Abisko | 531 ± 15 (25) | 66 ± 8.5 (18) | 3.39 ± 0.54 µm (22) | 4.2 ± 0.93 (13) | 1.22 ± 0.25 µm (12) |
| Stockholm | 514 ± 24 (25) | 45 ± 9.8 (19) | 2.87 ± 0.32 µm (21) | 3.8 ± 0.94 (12) | 1.16 ± 0.12 µm (10) |
| Wageningen | 463 ± 24 (25) | 43 ± 11.1 (11) | 2.73 ± 0.47 µm (25) | 2.2 ± 0.86 (15) | 0.87 ± 0.22 µm (15) |
| Jura | 489 ± 29 (21) | 42 ± 19.4 (16) | 2.86 ± 0.31 µm (22) | 2.1 ± 1.46 (7) | 1.09 ± 0.41 µm (7) |
| Callas | 482 ± 38 (24) | 34 ± 4.4 (13) | 3.63 ± 0.33 µm (25) | 2.0 ± 0.93 (8) | 0.94 ± 0.19 µm (8) |
| Pyrenees | 532 ± 25 (13) | 51 ± 17.2 (8) | 2.85 ± 0.49 µm (10) | 4.0 ± 0.81 (4) | 1.06 ± 0.08 µm (2) |
| Costa Brava | 488 ± 25 (19) | 48 ± 9.1 (9) | 3.82 ± 1.18 µm (19) | 2.4 ± 1.14 (5) | 0.95 ± 0.08 µm (5) |
| *P. napi* total average | 498 ± 35 (152) | 48 ± 13.9 (94) | 3.18 ± 0.70 µm (144) | 3.02 ± 1.34 (64) | 1.04 ± 0.26 µm (59) |

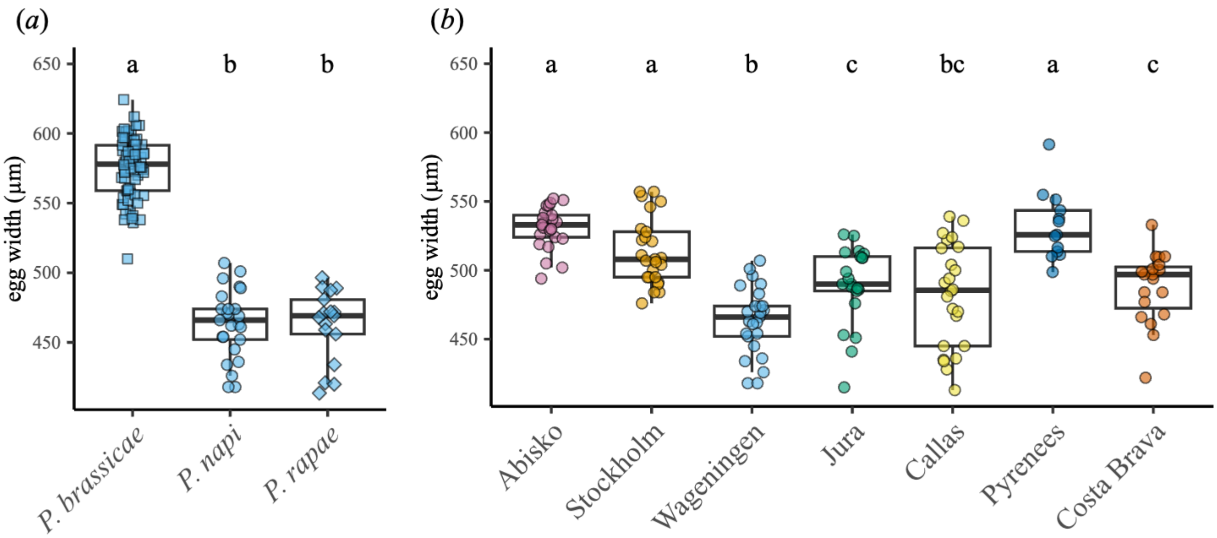


**Figure S2. Comparison of egg width (*a*) interspecifically between Pieris spp. and (*b*) intraspecifically between P. napi populations.** Points represent individual eggs (n = 111 and n = 152 eggs, respectively; Table S2; Table S9). Different letters indicate significant differences (*post hoc* using EMMs, $p > 0.05$).

### 3 Details of the seven *P. napi* populations

**Table S3. Summarized conditions for each of the European populations**

| population | latitude | longitude | Collected from field |
|---|---|---|---|
| **Abisko** | 68.34761 | 18.82977 | 2023 |
| **Stockholm** | 59.36447 | 18.03115 | 2023 |
| **Wageningen** | 51.97919 | 5.685259 | 2022 |
| **Jura** | 46.55577 | 6.08496 | 2022 |
| **Callas** | 43.5903 | 6.531266 | 2022 |
| **Pyrenees** | 42.32283 | 1.064298 | 2023 |
| **Costa Brava** | 42.26495 | 3.069001 | 2022 |

| **temperature seasonality** | **maximum temperature** | **precipitation** | **growing days** | **host plant** | **local conditions at host plant site** |
|---|---|---|---|---|---|
| 864.4 | 18 | 369.4 | 59 | *Arabidopsis* spp. and *Barbarea* spp. | Any present host plants near buildings and roads |
| 751.4 | 24.8 | 503.5 | 136 | *Alaria* spp. and *Bunias* spp. | Obtained from a park, not exclusively near water |
| 576.4 | 25.5 | 824 | 179 | various crop plants | Obtained near the river Eng, where conditions were cool and moist |
| 651.8 | 22.8 | 1348.3 | 130 | *Cardamine heptaphylla* | Moist and cool rocky ledges in a forested area |
| 601.7 | 27.8 | 829.4 | 202 | *Alaria petiolata* | Host plants in a relatively hot and dry area |
| 650.2 | 28.3 | 812.6 | 189 | *NA* | A dry and warm mountain regio with salty soil and little vegetation, where ovipositioning only occurred near water sources |
| 592.4 | 30.5 | 567.7 | 275 | *Lepidium draba* | Obtained from host plants that were shaded by trees in a dry agricultural area |

### 4 Correlation plots between egg traits

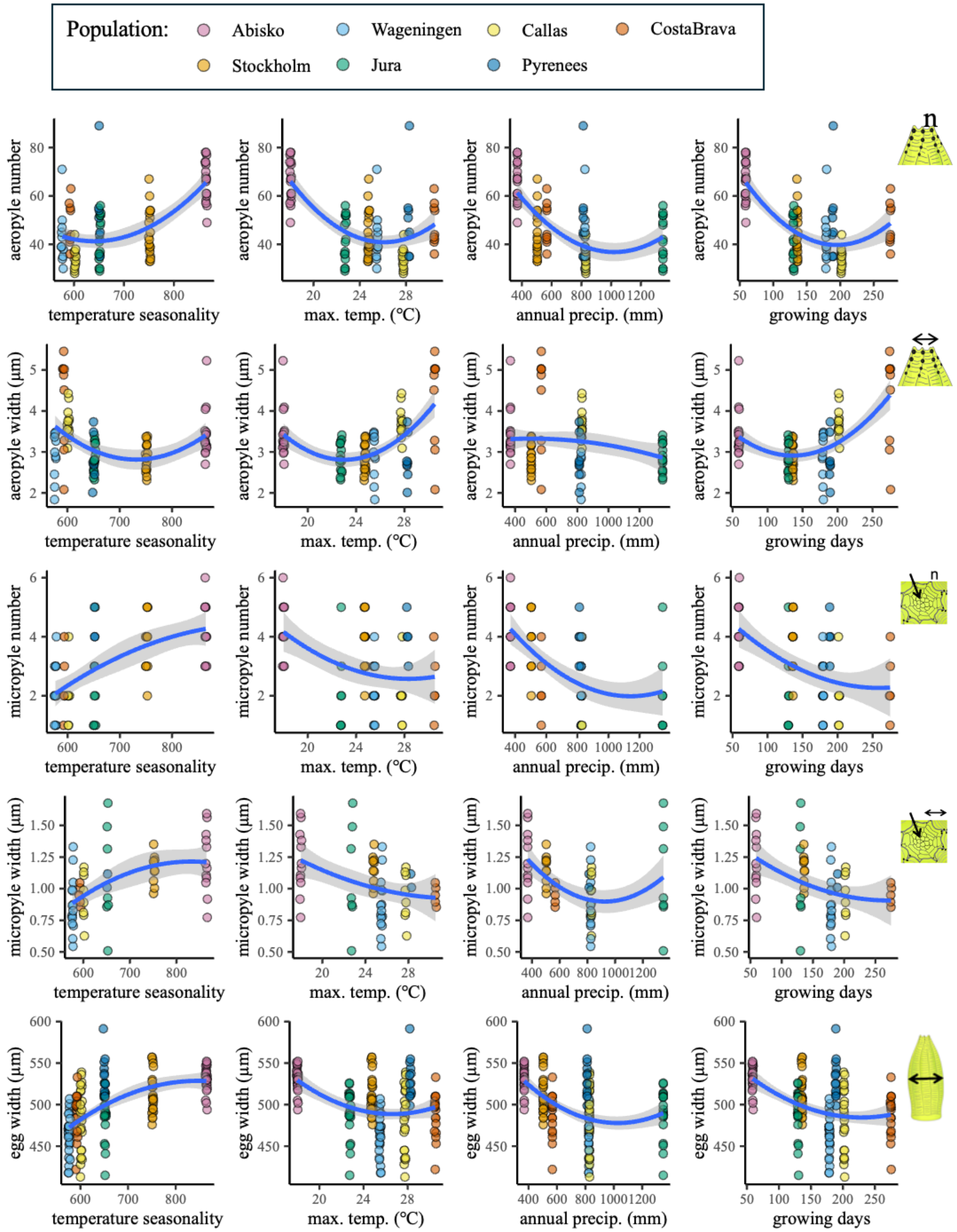


**Figure S3. Relations between climatic variables and egg pore traits.** (aeropyle number: n = 61 eggs, aeropyle width: n = 2830 aeropyles, micropyle number: n = 81 eggs, micropyle width: n = 255 micropyles; Table S2).

### 5 Piecewise Structural Equation Model (pSEM) selection

Our initial model included every possible association between egg traits and climatic variables (20 associations), as well as every possible association between egg trait to egg trait (nine associations, Figure S4). Plotting the data indicated the possibility of a polynomial relation between aeropyle traits and maximum temperature. We manually calculated the orthogonal polynomial of maximum temperature[89] and included this as an explanatory variable for aeropyle width and number (two associations).

The directionality of associations between egg traits were based on assumed biological relevance. The associations between climatic variables were incorporated in the model as random effects (six associations possible). To select the most parsimonious model that best explains the relationship among variables, we followed the procedure outlined in Boisseau and Woods, 2024[20]. In the first phase of model selection, individual associations were dropped step by step, testing which removed association led to the greatest reduction in the C statistic Information Criterion (CIC)[90], equivalent to d-sep determined Akaike Information Criterion or AIC from the current model. The current model is considered the best model if further dropping associations no longer leads to a decrease in CIC (see Figure S4). In the second phase of model selection, we tested alternative hypotheses on the directionality, and thus causal relation, of associations. Model fit was considered equivalent if $\Delta CIC \leq 2$, in which case the equivalent models should be averaged. Naturally, the causal relation between egg traits and climate is only in one direction. We further argued that the causal relation between egg width and micropyle number is one-way, as egg size is related by ecological and female traits[20,38]. There was only one alternative directionality possible after our model phase one: the pathway between the micropyle number and aeropyle number (Figure S5). As we used transposed datasets for aeropyle and micropyle width (every egg-pore value having its own data point), the directionality of their associations could not be reversed. Of the reduced model from phase one, only two alternative causal hypotheses could be tested (Figure S5). The alternative causal hypothesis did not improve model fit, Figure S5. Therefore, the reduced model (Figure S4*b*; Figure S5*a*), was selected as the final model. The model summary can be found in Table S4.

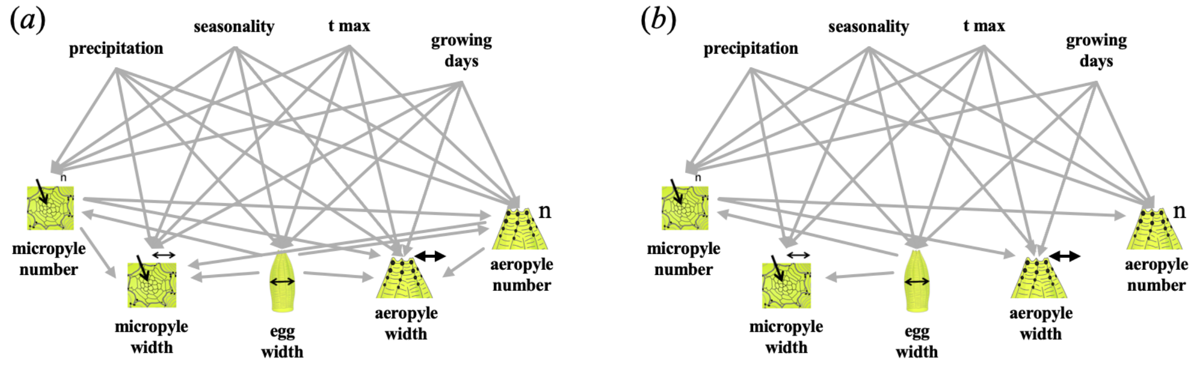

**Figure S4. Phase one of model selection, model reduction.** In the first phase, the **initial structural model (*a*)** including all possible associations was reduced step by step by dropping the association that led to the greatest increase in model fit (reduction in C statistic Information Criterion or CIC). Both polynomial relations (between both aeropyle traits and maximum temperature) did not contribute to model fit. The first phase resulted in a **reduced structural model (*b*)**. The initial model had a CIC of 97.42, and the reduced model had a CIC of 73.16

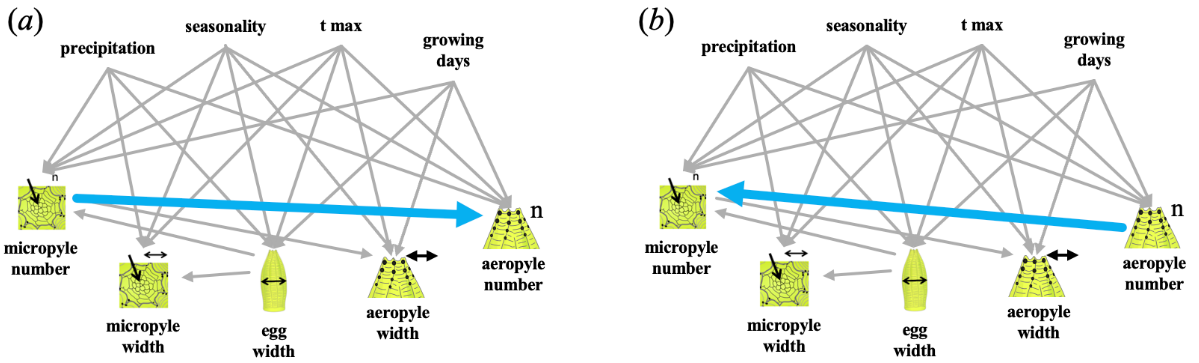


**Figure S5. Phase two of model selection testing alternative causal hypotheses.** In the second phase, the directionality of the pathway between the micropyle number and aeropyle number of the **reduced structural model (*a*)** was reversed to an **alternative structural model (*b*)**. This reversion did not lead to improved model fit. The C statistic Information Criterion (CIC) of the reduced model = 73.16, and the CIC of the alternative model = 84.94. ΔCIC > 2 and thus the reduced model (*a*) was selected as final model.

**Table S4. Standard estimates, standard errors (se), degrees of freedom (df), critical values (Crit.Value) and p-values for the final selected piecewise-SEM.**

| association | standard estimate | se | df | Crit.Value | p |
|---|---|---|---|---|---|
| **egg width <-- t. seasonality** | **1.187** | **0.203** | **147** | **5.849** | **<.001** |
| **egg width <-- t. max.** | **0.762** | **0.257** | **147** | **2.970** | **0.004** |
| **egg width <-- precipitation** | **0.231** | **0.111** | **147** | **2.083** | **0.039** |
| egg width <-- growing days | -0.149 | 0.312 | 147 | -0.477 | 0.634 |
| micropyle number <-- t. seasonality | 0.327 | 0.314 | 57 | 1.044 | 0.301 |
| micropyle number <-- t. max. | 0.565 | 0.422 | 57 | 1.341 | 0.185 |
| micropyle number <- precipitation | -0.198 | 0.172 | 57 | -1.154 | 0.253 |
| micropyle number <-- growing days | -0.569 | 0.531 | 57 | -1.071 | 0.289 |
| micropyle number <-- egg width | 0.199 | 0.119 | 57 | 1.672 | 0.100 |
| **aeropyle number <-- t. seasonality** | **0.779** | **0.367** | **33** | **2.120** | **0.042** |

| | | | | | |
|---|---|---|---|---|---|
| **aeropyle number <-- t. max** | **-1.572** | **0.380** | **33** | **-4.139** | **<.001** |
| aeropyle number <-- precipitation | -0.117 | 0.205 | 33 | -0.570 | 0.573 |
| **aeropyle number <-- growing days** | 1.853 | 0.512 | 33 | 3.618 | 0.001 |
| aeropyle number <- micropyle number | 0.225 | 0.146 | 33 | 1.541 | 0.133 |
| **aeropyle width <-- seasonality** | **0.685** | **0.145** | **36.40** | **22.274** | **<.001** |
| **aeropyle width <-- t. max** | **-1.182** | **0.223** | **35.59** | **28.216** | **<.001** |
| **aeropyle width <-- growing days** | **1.935** | **0.261** | **35.05** | **54.964** | **<.001** |
| aeropyle width <-- micropyle number | -0.135 | 0.079 | 36.61 | 2.899 | 0.097 |
| **micropyle width <-- t. seasonality** | **0.791** | **0.256** | **56.24** | **9.482** | **0.003** |
| micropyle width <-- t. max | 0.265 | 0.203 | 50.89 | 1.691 | 0.199 |
| micropyle width <-- precipitation | 0.227 | 0.155 | 61.23 | 2.130 | 0.150 |
| micropyle width <-- egg width | -0.135 | 0.120 | 63.93 | 1.269 | 0.264 |

### 6 Pairwise comparisons for population egg traits

**Table S5. Aeropyle number compared between the seven *Pieris napi* populations.** *Post hoc* using EMMs. Abbreviations: pop) population, est.) estimate, n) number, SE) standard error, df) degrees of freedom, t) t score, p) probability. ($\alpha$: $p > 0.05$).

| pop | pop | est. (n) | SE | df | t | p |
|---|---|---|---|---|---|---|
| **Abisko** | **Callas** | **32.18** | **3.58** | 87 | 8.98 | **<.001** |
| **Abisko** | **CostaBrava** | **18.00** | **4.02** | 87 | 4.48 | **<.001** |
| **Abisko** | **Jura** | **24.02** | **3.38** | 87 | 7.10 | **<.001** |
| **Abisko** | **Pyrenees** | **15.33** | **4.19** | 87 | 3.66 | **0.008** |
| **Abisko** | **Stockholm** | **21.81** | **3.24** | 87 | 6.73 | **<.001** |
| **Abisko** | **Wageningen** | **23.52** | **3.77** | 87 | 6.24 | **<.001** |
| **Callas** | **CostaBrava** | **-14.18** | **4.27** | 87 | -3.32 | **0.022** |
| Callas | Jura | -8.16 | 3.68 | 87 | -2.22 | 0.297 |
| **Callas** | **Pyrenees** | **-16.85** | **4.43** | 87 | -3.81 | **0.005** |
| Callas | Stockholm | -10.37 | 3.55 | 87 | -2.93 | 0.064 |
| Callas | Wageningen | -8.66 | 4.03 | 87 | -2.15 | 0.335 |
| CostaBrava | Jura | 6.02 | 4.10 | 87 | 1.47 | 0.763 |
| CostaBrava | Pyrenees | -2.67 | 4.79 | 87 | -0.56 | 0.998 |
| CostaBrava | Stockholm | 3.81 | 3.99 | 87 | 0.96 | 0.962 |
| CostaBrava | Wageningen | 5.52 | 4.43 | 87 | 1.25 | 0.874 |
| Jura | Pyrenees | -8.69 | 4.26 | 87 | -2.04 | 0.399 |
| Jura | Stockholm | -2.21 | 3.34 | 87 | -0.66 | 0.994 |
| Jura | Wageningen | -0.51 | 3.86 | 87 | -0.13 | 1.000 |

| | | | | | | |
|---|---|---|---|---|---|---|
| Pyrenees | Stockholm | 6.47 | 4.15 | 87 | 1.56 | 0.708 |
| Pyrenees | Wageningen | 8.18 | 4.58 | 87 | 1.79 | 0.560 |
| Stockholm | Wageningen | 1.71 | 3.73 | 87 | 0.46 | 0.999 |

**Table S6. Aeropyle width compared between the seven *Pieris napi* populations.** *Post hoc* using EMMs. Abbreviations: pop) population, est.) estimate, n) number, SE) standard error, df) degrees of freedom, t) t score, p) probability. ($\alpha$: $p > 0.05$).

| **pop** | **pop** | **est. (µm)** | **SE** | **df** | **t** | **p** |
|---|---|---|---|---|---|---|
| Abisko | Callas | -0.287 | 0.203 | 88.8 | -1.42 | 0.792 |
| **Abisko** | **CostaBrava** | **-0.820** | **0.226** | 85.0 | -3.63 | **0.008** |
| Abisko | Jura | 0.558 | 0.191 | 87.1 | 2.93 | 0.064 |
| Abisko | Pyrenees | 0.609 | 0.235 | 85.1 | 2.59 | 0.143 |
| **Abisko** | **Stockholm** | **0.579** | **0.181** | 85.5 | 3.20 | **0.031** |
| Abisko | Wageningen | 0.509 | 0.212 | 86.5 | 2.40 | 0.212 |
| Callas | CostaBrava | -0.533 | 0.245 | 89.1 | -2.18 | 0.318 |
| **Callas** | **Jura** | **0.845** | **0.213** | 92.2 | 3.97 | **0.003** |
| **Callas** | **Pyrenees** | **0.896** | **0.253** | 88.8 | 3.54 | **0.011** |
| **Callas** | **Stockholm** | **0.866** | **0.204** | 91.3 | 4.24 | **0.001** |
| **Callas** | **Wageningen** | **0.796** | **0.232** | 90.8 | 3.43 | **0.016** |
| **CostaBrava** | **Jura** | **1.378** | **0.235** | 88.0 | 5.87 | **<.001** |
| **CostaBrava** | **Pyrenees** | **1.429** | **0.272** | 86.2 | 5.25 | **<.001** |
| **CostaBrava** | **Stockholm** | **1.399** | **0.227** | 87.0 | 6.16 | **<.001** |
| **CostaBrava** | **Wageningen** | **1.329** | **0.252** | 87.4 | 5.26 | **<.001** |
| Jura | Pyrenees | 0.051 | 0.244 | 87.7 | 0.21 | 1.000 |
| Jura | Stockholm | 0.022 | 0.192 | 89.9 | 0.11 | 1.000 |
| Jura | Wageningen | -0.049 | 0.222 | 89.7 | -0.22 | 1.000 |
| Pyrenees | Stockholm | -0.029 | 0.236 | 86.9 | -0.12 | 1.000 |
| Pyrenees | Wageningen | -0.100 | 0.261 | 87.3 | -0.38 | 1.000 |
| Stockholm | Wageningen | -0.070 | 0.214 | 88.8 | -0.33 | 1.000 |

**Table S7. Micropyle number compared between the seven *Pieris napi* populations.** *Post hoc* using EMMs. Abbreviations: pop) population, est.) estimate, n) number, SE) standard error, df) degrees of freedom, t) t score, p) probability. ($\alpha$: $p > 0.05$).

| **pop** | **pop** | **est. (n)** | **SE** | **df** | **t** | **p** |
|---|---|---|---|---|---|---|
| **Abisko** | **Callas** | **2.23** | **0.45** | 57 | 4.99 | **<.001** |
| **Abisko** | **CostaBrava** | **1.83** | **0.52** | 57 | 3.50 | **0.015** |

| | | | | | | |
|---|---|---|---|---|---|---|
| **Abisko** | **Jura** | **2.09** | **0.47** | 57 | 4.48 | **<.001** |
| Abisko | Pyrenees | 0.23 | 0.57 | 57 | 0.41 | 1.000 |
| Abisko | Stockholm | 0.40 | 0.40 | 57 | 1.00 | 0.953 |
| **Abisko** | **Wageningen** | **2.03** | **0.38** | 57 | 5.39 | **<.001** |
| Callas | CostaBrava | -0.40 | 0.57 | 57 | -0.71 | 0.992 |
| Callas | Jura | -0.14 | 0.52 | 57 | -0.28 | 1.000 |
| **Callas** | **Pyrenees** | **-2.00** | **0.61** | 57 | -3.28 | **0.028** |
| **Callas** | **Stockholm** | **-1.83** | **0.45** | 57 | -4.04 | **0.003** |
| Callas | Wageningen | -0.20 | 0.44 | 57 | -0.46 | 0.999 |
| CostaBrava | Jura | 0.26 | 0.58 | 57 | 0.44 | 0.999 |
| CostaBrava | Pyrenees | -1.60 | 0.67 | 57 | -2.40 | 0.219 |
| CostaBrava | Stockholm | -1.43 | 0.53 | 57 | -2.71 | 0.115 |
| CostaBrava | Wageningen | 0.20 | 0.51 | 57 | 0.39 | 1.000 |
| Jura | Pyrenees | -1.86 | 0.62 | 57 | -2.98 | 0.061 |
| **Jura** | **Stockholm** | **-1.69** | **0.47** | 57 | -3.57 | **0.012** |
| Jura | Wageningen | -0.06 | 0.46 | 57 | -0.13 | 1.000 |
| Pyrenees | Stockholm | 0.17 | 0.57 | 57 | 0.29 | 1.000 |
| **Pyrenees** | **Wageningen** | **1.80** | **0.56** | 57 | 3.22 | **0.033** |
| **Stockholm** | **Wageningen** | **1.63** | **0.39** | 57 | 4.24 | **0.002** |

**Table S8. Micropyle width compared between the seven *Pieris napi* populations.** *Post hoc* using EMMs. Abbreviations: pop) population, est.) estimate, n) number, SE) standard error, df) degrees of freedom, t) t score, p) probability. ($\alpha$: $p > 0.05$).

| **pop** | **pop** | **est. (µm)** | **SE** | **df** | **t** | **p** |
|---|---|---|---|---|---|---|
| Abisko | Jura | 0.121 | 0.106 | 53.3 | 1.14 | 0.913 |
| Abisko | Callas | 0.278 | 0.102 | 56.7 | 2.72 | 0.113 |
| Abisko | CostaBrava | 0.271 | 0.116 | 50.4 | 2.35 | 0.244 |
| Abisko | Pyrenees | 0.150 | 0.154 | 40.2 | 0.97 | 0.958 |
| Abisko | Stockholm | 0.063 | 0.086 | 38.6 | 0.73 | 0.990 |
| **Abisko** | **Wageningen** | **0.330** | **0.082** | 47.5 | 4.00 | **0.004** |
| Jura | Callas | 0.157 | 0.123 | 65.4 | 1.28 | 0.859 |
| Jura | CostaBrava | 0.151 | 0.134 | 58.3 | 1.12 | 0.919 |
| Jura | Pyrenees | 0.029 | 0.169 | 45.4 | 0.17 | 1.000 |
| Jura | Stockholm | -0.058 | 0.110 | 52.7 | -0.53 | 0.998 |
| Jura | Wageningen | 0.209 | 0.107 | 61.4 | 1.96 | 0.453 |
| Callas | CostaBrava | -0.007 | 0.131 | 60.9 | -0.05 | 1.000 |
| Callas | Pyrenees | -0.128 | 0.166 | 46.3 | -0.77 | 0.987 |

| | | | | | | |
|---|---|---|---|---|---|---|
| Callas | Stockholm | -0.215 | 0.106 | 55.8 | -2.03 | 0.407 |
| Callas | Wageningen | 0.052 | 0.103 | 66.0 | 0.50 | 0.999 |
| CostaBrava | Pyrenees | -0.122 | 0.175 | 44.9 | -0.70 | 0.992 |
| CostaBrava | Stockholm | -0.209 | 0.119 | 50.1 | -1.76 | 0.584 |
| CostaBrava | Wageningen | 0.058 | 0.116 | 56.6 | 0.50 | 0.999 |
| Pyrenees | Stockholm | -0.087 | 0.157 | 40.3 | -0.55 | 0.998 |
| Pyrenees | Wageningen | 0.180 | 0.155 | 42.8 | 1.16 | 0.904 |
| Stockholm | Wageningen | 0.267 | 0.087 | 47.3 | 3.08 | 0.050 |

**Table S9. Egg width compared between the seven *Pieris napi* populations.** *Post hoc* using EMMs. Abbreviations: pop) population, est.) estimate, n) number, SE) standard error, df) degrees of freedom, t) t score, p) probability. ($\alpha$: $p > 0.05$).

| **pop** | **pop** | **est. (µm)** | **SE** | **df** | **t** | **p** |
|---|---|---|---|---|---|---|
| **Abisko** | **Callas** | **48.8** | **7.6** | **145** | **6.45** | **<0.001** |
| **Abisko** | **CostaBrava** | **42.9** | **8.1** | **145** | **5.32** | **<0.001** |
| **Abisko** | **Jura** | **41.6** | **7.9** | **145** | **5.30** | **<0.001** |
| Abisko | Pyrenees | -1.1 | 9.1 | 145 | -0.12 | 1.000 |
| Abisko | Stockholm | 17.0 | 7.5 | 145 | 2.27 | 0.265 |
| **Abisko** | **Wageningen** | **67.3** | **7.5** | **145** | **8.98** | **<0.001** |
| Callas | CostaBrava | -5.9 | 8.1 | 145 | -0.73 | 0.991 |
| Callas | Jura | -7.2 | 7.9 | 145 | -0.91 | 0.970 |
| **Callas** | **Pyrenees** | **-50.0** | **9.1** | **145** | **-5.47** | **<0.001** |
| **Callas** | **Stockholm** | **-31.8** | **7.6** | **145** | **-4.20** | **<0.001** |
| Callas | Wageningen | 18.5 | 7.6 | 145 | 2.44 | 0.190 |
| CostaBrava | Jura | -1.3 | 8.4 | 145 | -0.16 | 1.000 |
| **CostaBrava** | **Pyrenees** | **-44.0** | **9.5** | **145** | **-4.61** | **<0.001** |
| **CostaBrava** | **Stockholm** | **-25.9** | **8.1** | **145** | **-3.21** | **0.027** |
| **CostaBrava** | **Wageningen** | **24.4** | **8.1** | **145** | **3.02** | **0.046** |
| **Jura** | **Pyrenees** | **-42.7** | **9.4** | **145** | **-4.57** | **<0.001** |
| **Jura** | **Stockholm** | **-24.6** | **7.9** | **145** | **-3.13** | **0.034** |
| **Jura** | **Wageningen** | **25.7** | **7.9** | **145** | **3.28** | **0.022** |
| Pyrenees | Stockholm | 18.2 | 9.1 | 145 | 2.00 | 0.417 |
| **Pyrenees** | **Wageningen** | **68.4** | **9.1** | **145** | **7.55** | **<0.001** |
| **Stockholm** | **Wageningen** | **50.3** | **7.5** | **145** | **6.71** | **<0.001** |
